# Machine Learning-Guided Discovery of Bacterial-Selective Membrane-Active Compounds Reveals Mechanistic Bias in Antibiotic Training Datasets

**DOI:** 10.64898/2026.06.08.730938

**Authors:** Connor Chain, Soodabeh Ghaffari, Syrine Belakaria, Joseph P. Sheehan, Ido Irani, Chi-Yun Wu, Hahn Kim, Barbara E Engelhardt, Zemer Gitai

## Abstract

The rise of antibiotic resistance necessitates the discovery of antibacterial compounds with novel mechanisms of action (MoAs). Recent machine learning approaches have shown promise in antibacterial compound discovery, but often identify derivatives of known antibiotic classes rather than mechanistically novel compounds. Previous approaches applied Tanimoto similarity filters at the end of screening pipelines, but this method has substantial drawbacks: Tanimoto similarity can be misleading in chemical space, and post-hoc filtering does not influence what activity models learn to prioritize. Here, we present a machine learning pipeline that addresses chemical novelty upfront by employing an XGBoost-based MoA classifier to explicitly prioritize compounds predicted to have mechanisms distinct from known antibiotic classes, combined with graph neural networks for antibacterial activity and toxicity prediction. Applied to the Zinc20 database, our approach successfully identified non-toxic antibacterial compounds structurally distinct from known antibiotics. Notably, the majority of these hits exhibited membrane-targeting activity with selectivity for bacterial cells over mammalian cells, suggesting potential for next-generation membrane-active antibiotics. However, we did not identify compounds with novel protein targets. Systematic analysis revealed that this limitation stems from mechanistic bias in training data rather than model architecture. Specifically, our activity model learned to preferentially score compounds similar to specific groups in the training data, thus overrepresenting certain MoA classes including membrane-active compounds. Even substantial model architecture and training data enhancements did not overcome this constraint. Our findings demonstrate that the primary bottleneck for discovering mechanistically novel antibiotics is the scarcity of diverse, mechanistically-annotated training data. This work provides both a methodological framework for mechanism-aware screening and critical insights into data requirements for genuinely novel antibiotic discovery.

## Introduction

The prevalence of antibiotic-resistant infections is increasing at an alarming rate. In 2019, 1.27 million deaths could be directly attributed to antimicrobial-resistant (AMR) infections around the world, with another 4.95 million deaths associated with AMR infections [1], where resistance was present and likely contributed to worse outcomes, but was not necessarily the sole cause of death. These infections are predicted to skyrocket by 2050, with deaths from AMR infections expected to exceed those from cancer [2]. Clinical isolates of *A. baumannii* are already capable of resisting all available antibiotics [3]. Over the past two decades, clinical isolates harboring newly evolved resistance mechanisms, including carbapenemases and colistin resistance determinants. have rapidly emerged and disseminated across diverse bacterial species [4]. This situation is alarming.

To address this threat to public health, it is imperative to identify new antibiotics capable of overcoming the existing resistance to antibiotics in use. Molecules with novel mechanisms of action (MoAs) are particularly valuable as they are less susceptible to the same resistance mechanisms that persist in newly arising antibiotic-resistant isolates. Historically, antibiotics have often been found by screening natural products to identify compounds in the environment capable of inhibiting bacterial growth [5, 6]. However, this strategy has produced diminishing returns in recent years, often leading to the rediscovery of already existing classes of antibiotics [7, 8].

An alternative strategy for identifying antimicrobial compounds is to screen large libraries of small synthetic molecules. However, these efforts can be challenging for several reasons. First, screening such large libraries can be time-consuming and expensive [9]. Second, the selection of different screen outputs or chemical library inputs can strongly bias screen results. Antibiotic screening efforts have largely prioritized bacterial killing (antibacterial activity). But additional factors like low toxicity, desirable drug-like properties, and novel MoA are also important considerations for how to prioritize candidates. Due to these limitations, the screening of small synthetic libraries has often led to promising initial hits but relatively few clinically useful antibiotics, leading to pharmaceutical companies abandoning the field and defunding many of their antibiotic discovery programs [10, 11].

Recently, *in silico* strategies have been developed to improve drug discovery [12–14] and increase the capacity and effectiveness of small molecule library screens. These strategies involve experimentally testing relatively small libraries of compounds and using the data collected to train machine learning (ML) models. ML models can predict desirable outcomes for new molecular collections in silico, prioritizing promising candidate molecules for experimental validation.

With the recent increased interest in accelerating drug design using machine learning, several packages have been developed to facilitate implementing ML pipelines for molecular design. Of particular interest are the DeepChem [12] and Chemprop [15, 16] packages, which allow graph neural network models to predict chemical characteristics of small molecules. Chemprop uses a directed message-passing neural network (D-MPNN) that propagates information along bonds in a molecular graph, enabling the model to capture local chemical patterns formed by atoms and their connecting bonds, making it well-suited for predicting molecular properties.

Recent work leveraging the Chemprop library [15] identified the novel antibiotic compound halicin [17], and has since been used in subsequent studies to identify small molecules capable of killing stationary phase bacteria and *A. baumaanni* [18, 19]. More recent efforts in ML antibiotic discovery have also incorporated toxicity predictions within their models [20]. While these approaches have contributed substantially to the field, they have limitations that need to be overcome to reach their full potential.

A central challenge in antibiotic discovery is identifying compounds with genuine chemical and mechanistic novelty: molecules that are structurally distinct from existing antibiotics and therefore less likely to be subject to cross-resistance. Previous ML-based approaches have attempted to address this using Tanimoto similarity as a post-hoc filter at the end of the screening pipeline [17]. However, Tanimoto similarity is not an appropriate metric to use, particularly in small molecule space [21] where, for example, slight perturbations to structures containing rings can skew the Tanimoto similarity of two molecules to appear more different than they actually are. Moreover, applying novelty filters only at the final selection stage leads to an inefficient resource expenditure, requiring the (often prohibitively costly) acquisition of many candidate compounds that could inhibit bacteria through known mechanisms.

The data used for training ML models is also an important factor to consider as it will directly impact the outcome of the method. Using training data containing classes of known antibiotics will naturally lead to predictions that yield more compounds with high antibacterial activity and low toxicity; however, most of these compounds represent derivatives of known antibiotics and consequently restrict the exploration of novel compounds with alternative mechanisms of action in chemical space. Additionally, training data are often imbalanced, with many more compounds lacking antibiotic activity (negative labels) than compounds with antibiotic activity (positive labels). The training data are also limited by compounds that have already been tested for antibiotic activity and toxicity experimentally, which is a small slice of the vastness of chemical space. Moreover, the test set of uncharacterized compounds to label in this in silico screen is often limited to a specific type of compound, and experimental validation of the top predictions is strictly limited by experimental resources.

Together, these limitations lead to the problem that newly found compounds in recent work [18, 19], which are initially described as novel, actually share structural motifs and consequently, mechanisms of action with known antibiotics. For example, many of the compounds identified in Stokes et al. 2020 [17] have motifs reminiscent of quinolone antibiotics. Other studies have overstated the effectiveness of ML approaches. For example, the compound Semapimod did not require Chemprop to be discovered, since this compound was identified in the original screen used as training data [19]. Truly mechanistically novel antibiotics remain elusive, providing room for innovation in the machine-learning antibiotic discovery field.

In this work, we present two machine learning approaches that address mechanistic novelty upfront rather than as an afterthought. Unlike previous methods that apply Tanimoto-based filters at the end of the pipeline, we employ a mechanism-of-action (MoA) classifier at the beginning of our workflow to explicitly prioritize compounds predicted to have novel mechanisms. Our approach recognizes that the goal of novelty in antibiotic discovery is not merely abstract structural dissimilarity, but rather the identification of molecules with distinct chemical types that are unlikely to be subject to existing resistance mechanisms. Both methods leverage graph neural networks (GNNs) to predict antibiotic activity and toxicity, while an extreme gradient boosting (XGBoost) model evaluates mechanistic novelty by classifying molecules based on their dissimilarity to known antibiotic classes.

In the first approach, we leverage two general graph convolutional neural network (GCN) models introduced by Duvenaud et al. (2015) [22] and implemented within the DeepChem package [12]. These models are combined to filter datasets of new molecules by sequentially applying thresholds across the predictions of activity, toxicity, and MoA novelty. The compounds are subsequently filtered using quality filters of the physicochemical properties of desirable drugs. Then, candidates are validated experimentally for antibiotic activity and toxicity. In the second approach, we use two directed message-passing neural network (D-MPNN) models from the Chemprop package [15] to predict activity and toxicity. This approach expands the XGBoost model into a larger ensemble, improving its capacity to manage data imbalance and to incorporate predictions across additional classes. The resulting models are integrated to filter datasets of new molecules by solving a multi-objective optimization problem, balancing the outputs of the three models, and selecting compounds that achieve the best Pareto-front tradeoff.

When we applied these workflows to screen for antibiotic compounds in the Zinc20 [23] database, we successfully identified and validated compounds with antibacterial activity that were nontoxic, met our thresholds for favorable drug properties, and exhibited structural dissimilarities to known antibiotics by our MoA classifier. Deeper investigation revealed that many of these compounds are membrane-active yet non-toxic. These findings raise the possibility that bacterial-selective membrane disruption could be a powerful yet under-utilized approach to developing novel antibiotics. More broadly, our results provide valuable insights into the power and limitations of structure-based ML approaches and reveal an important bias in current training datasets toward membrane-active compounds, highlighting the need for more mechanistically diverse training data to enable the discovery of antibiotics with truly novel protein targets.

## Results

### Machine learning pipeline for antibiotic discovery with upfront novelty filtering

Our goal was to design a machine learning approach to predict the antibacterial activity (bacterial growth inhibition), toxicity (mammalian cell killing), and mechanistic novelty of compounds based on their molecular structures. To this end, we developed three separate models: two graph convolutional network (GCN) models using the DeepChem toolkit [12] for antibacterial activity and toxicity prediction, and an XGBoost model for mechanism-of-action (MoA) classification (Figure 1A).

**Fig 1.**
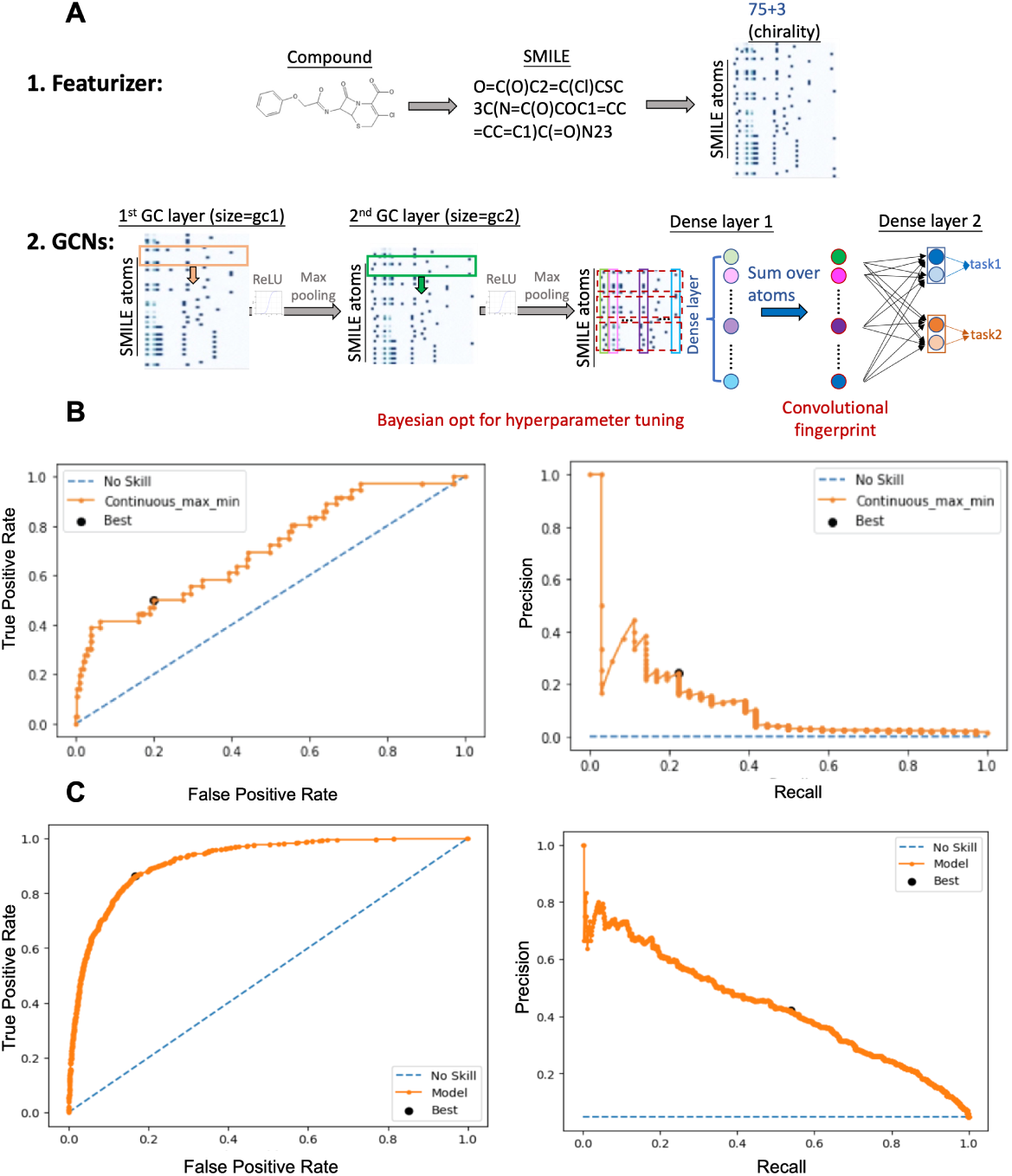
Deepchem pipeline for identifying novel antibiotics. (A) Overview of the DeepChem GCN architecture. Molecular features are derived from SMILES representations and processed through GCN layers for vectorization and downstream tasks (B) ROC and PR curve for the antibiotic activity model and (C) ROC and PR curve for the toxicity model.

#### Antibacterial activity model training and validation

To train the antibacterial activity model, we collected growth inhibition rates of small molecules by treating *E. coli* lptD4213 with 50*µ*M of each compound and measuring the optical density at 600nm (OD600) after 24 hours compared to an untreated control. *E. coli* lptD4213 [24] was chosen as this species has increased permeability compared to wild-type E. coli, allowing us to capture compounds with biological activity that otherwise would be unable to cross the outer membrane. To increase the size of our training data, we included growth inhibition values from the Shared Platform for Antibiotic Research and Knowledge [25] (SPARK) of *E. coli* ATCC 25922 treated with 32*µ*g/mL for a total of 122,851 compounds with growth inhibition values (Table 1). We also trained a classification-based GCN model by binarizing the values for each compound based on a growth inhibition threshold of 80%, classifying compounds as active if their inhibition exceeded this threshold. We found that the regression-based model had better performance than the classification model, so we used that model for subsequent analysis. For testing, we used a collection of 2,208 compounds with binary activity labels experimentally determined from a subset of small molecules from the Chembridge DIVERSet Library. To validate the DeepChem-based model for antibacterial activity, we split the labeled datasets randomly into 80% for training and 20% for validation, with each split containing the same percentage of active compounds. We divided the predicted antibiotic activity values by 100 to interpret them as probabilities and used classification metrics, including the area under the curve (AUC) for receiver operating characteristic (ROC) curves and precision-recall (PR) curves, to evaluate the model. For the antibiotic activity model, we achieved an ROC AUC of 0.71 and a PR AUC of 0.13 (Figure 1B).

**Table 1.**
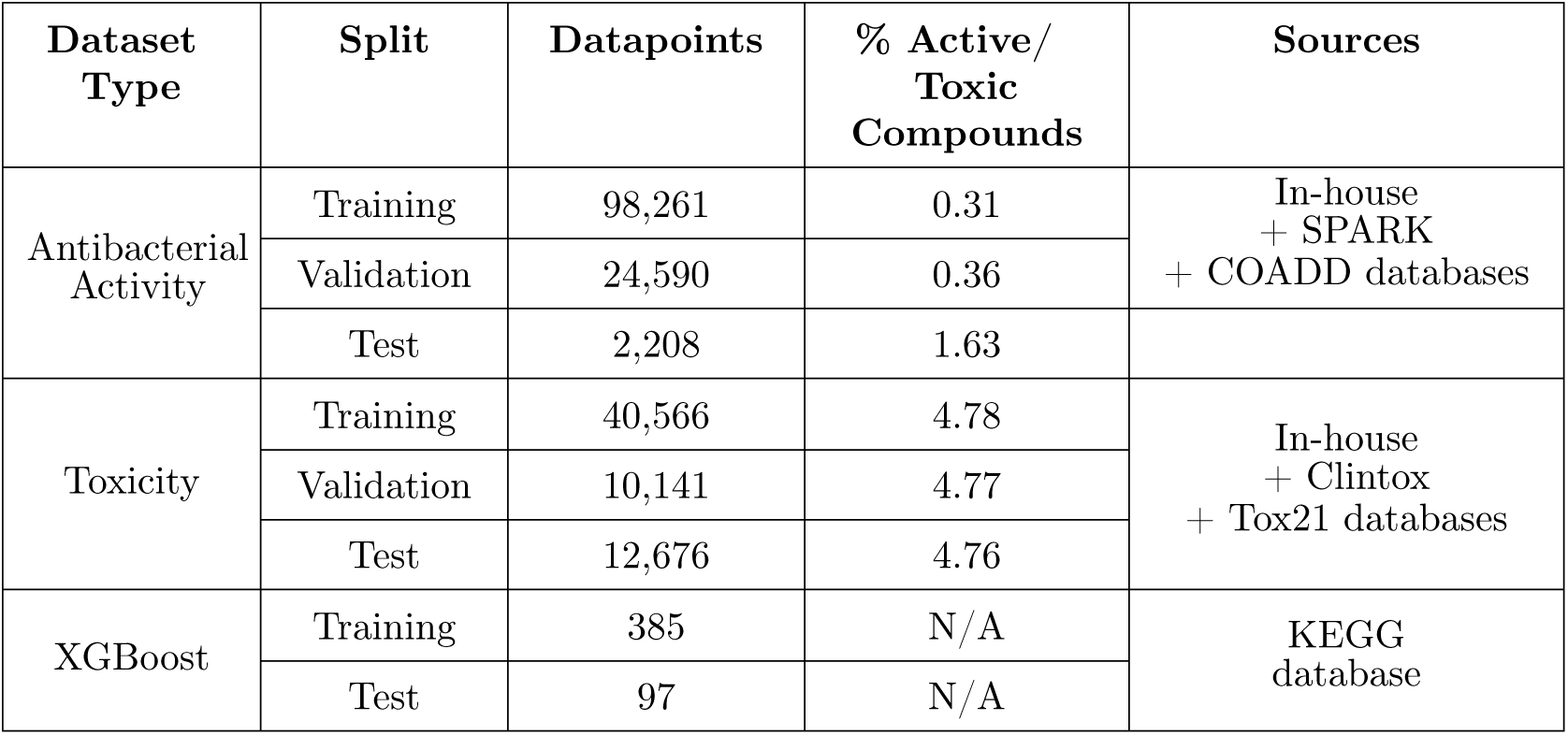
Summary of datasets used in the study for the DeepChem-based method.

| Dataset Type | Split | Datapoints | % Active/<br>Toxic<br>Compounds | Sources |
| --- | --- | --- | --- | --- |
| Antibacterial Activity | Training | 98,261 | 0.31 | In-house<br>+ SPARK<br>+ COADD databases |
|  | Validation | 24,590 | 0.36 |  |
|  | Test | 2,208 | 1.63 |  |
| Toxicity | Training | 40,566 | 4.78 | In-house<br>+ Clintox<br>+ Tox21 databases |
|  | Validation | 10,141 | 4.77 |  |
|  | Test | 12,676 | 4.76 |  |
| XGBoost | Training | 385 | N/A | KEGG<br>database |
|  | Test | 97 | N/A |  |

#### Mammalian cell toxicity model training and validation

To train the toxicity model, we collected toxicity data in an analogous study using the same small molecule library that treated *E. coli* lptD4213 to treat Huh7 cells at 50*µ*M [26]. Compounds in wells with *<* 50% cell coverage after four days in the presence of each small molecule were classified as toxic. To increase the size of our toxicity training data, we combined the binary toxicity information of 56,916 compounds curated in-house, 1,484 compounds (112 toxic) from the Clintox database [27], and 7,831 compounds (2,872 toxic) from the Tox21 database [28]. With the combined binary toxicity data, we trained a GCN classifier to predict if a new compound is toxic. For the toxicity model, the ROC AUC and PR AUC were 0.92 and 0.43, respectively (Figure 1C).

To fairly compare results from our models with those from Chemprop, we tested each of the activity and toxicity models on the same test set of molecules. The results showed that our model performed similarly to Chemprop (Table 2) when both models were trained and tested on the same data splits.

**Table 2.** Performance comparison of DeepChem and Chemprop models on the test data. ROC-AUC values for DeepChem and Chemprop trained and tested on the same dataset.

| Model | Test Data | ROC-AUC |
| --- | --- | --- |
| DeepChem | Antibiotic | 0.71 |
| DeepChem | Toxicity | 0.92 |
| Chemprop | Antibiotic | 0.68 |
| Chemprop | Toxicity | 0.93 |

#### MoA classifier for prioritizing mechanistic novelty

To prioritize molecules with novel mechanisms of action (MoAs), we trained an XGBoost model to classify molecules into known antibiotic types. The XGBoost model determines the probability of a molecule fitting into one of thirteen categories of known antibiotic targets based on its Morgan fingerprint. We used a list of known antibiotics and their targets from the Kyoto Encyclopedia of Genes and Genomes (KEGG) database [29] ombined with in-house labeled antibiotics as our dataset. We split the dataset into training and testing sets using an 80%–20% stratified split to maintain the distribution of the labels. To assess the performance of the trained XGBoost MoA classifier, we computed the precision, recall, and F1 score of the test dataset (Table 3). Using the testing dataset, we calculated an average precision of 93%, recall of 94% and F1 score of 91%, indicating that the classifier was working well (Figure 2).

**Table 3.**
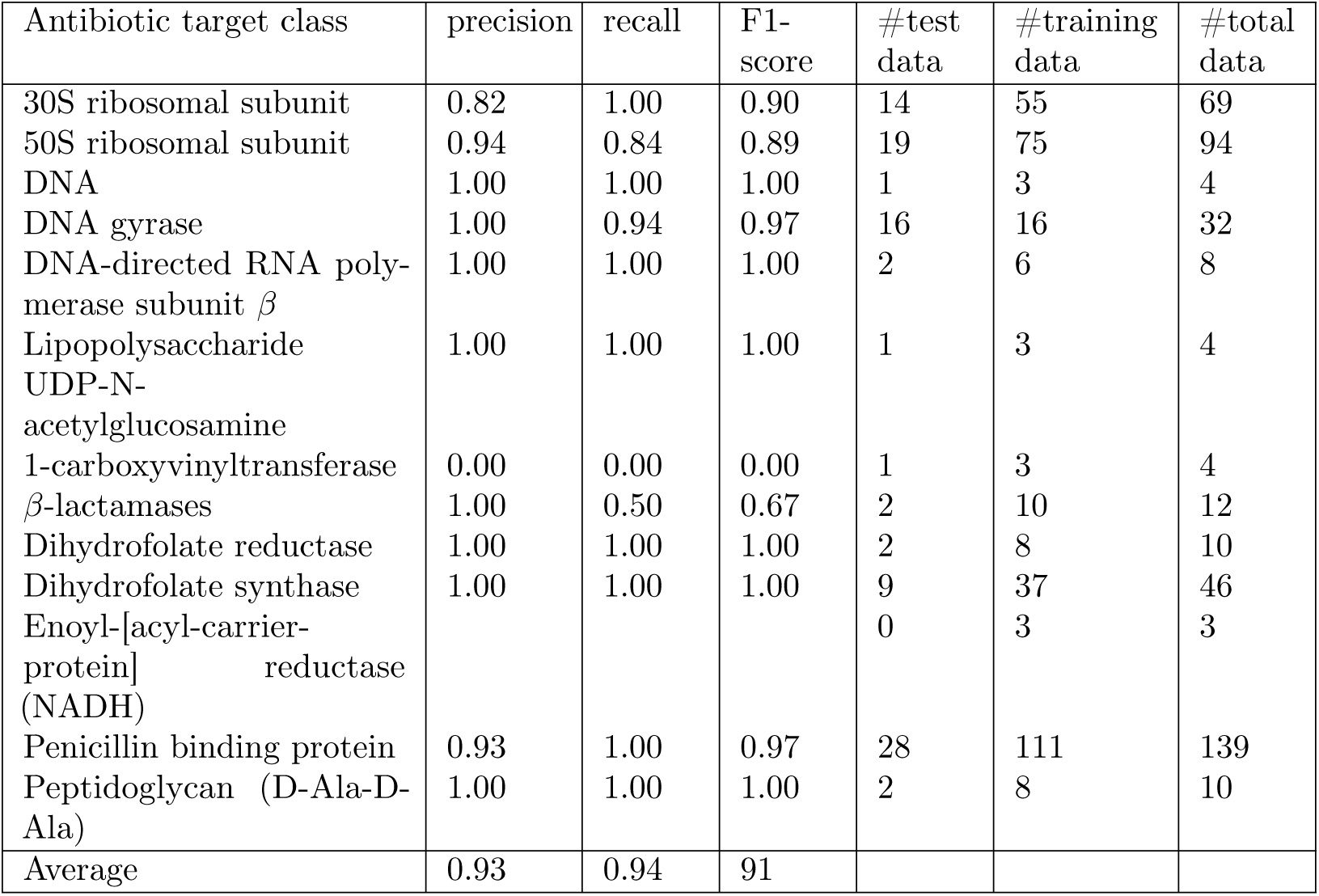
XGBoost classifier performance on 13 known antibiotic classes. Individual values are given for known antibiotics classes curated from the KEGG database. UDP-N-acetylglucosamine 1-carboxyvinyltransferase only had one representative molecule in the test data, which was misclassified.

**Fig 2.**
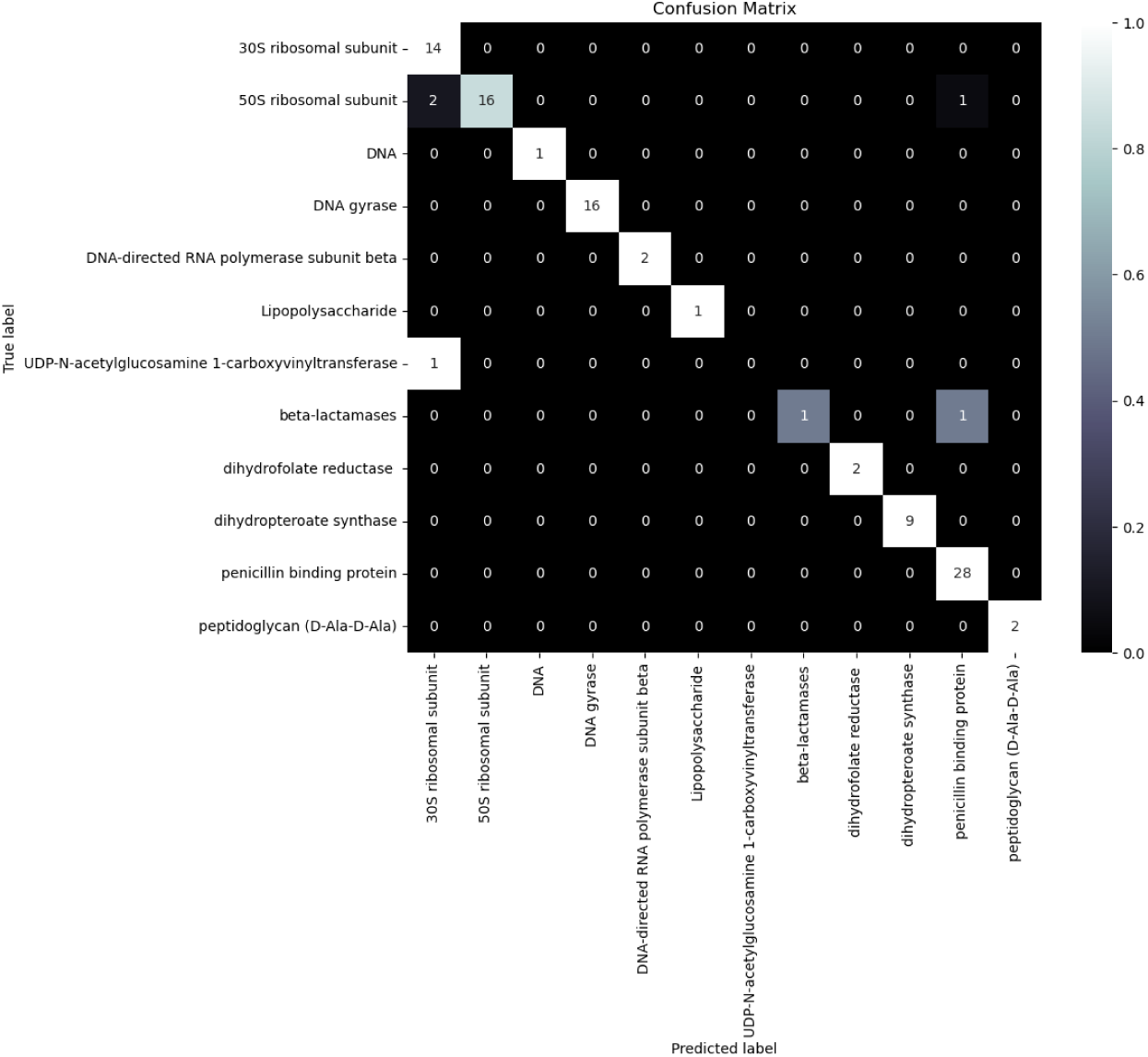
Confusion matrix of the XGBooost model for MoA classification. As Enoyl-[acyl-carrier-protein] reductase (NADH) does not have any data in test dataset, it is removed from the confusion matrix.

Our goal for the XGBoost model was to prioritize compounds that have novel mechanisms to kill bacteria (i.e., compounds that are functionally dissimilar to existing antibiotics). To identify molecules that have an MoA different from the 13 known classes using a model trained only on molecules in these classes, we used classification uncertainty to capture novel MoA. In particular, our multi-class classifier returns 13 different values capturing the probability that a new molecule *x* belongs to one of the 13 classes *c_i_*, *p*(*c_i_*|*x*), where *i* ∈ {1*, . . . ,* 13}. If all of the probabilities have low values, then the model is struggling to assign any of the known class labels to the molecule. To this end, we defined the dissimilarity score as 1 − max *p*(*c_i_*|*x*) to indicate the level of dissimilarity of new molecule *x* to the 13 known antibiotic MoA classes.

### ML-driven identification of candidate novel antibacterial compounds

To search for antimicrobial small molecules in silico, we applied our trained models to the Zinc20 database [23] (Figure 3A), which contains around 884 million compounds. Specifically, we applied an initial filter that led to a subset of approximately 11 million compounds that are both commercially available (allowing us to experimentally test them) and drug-like. These are molecules whose LogP values do not exceed 5 and whose molecular weights range from 250 to 500 Daltons based on Lipinski’s Rule of Five [30]. We then applied our three machine learning models (Activity, Toxicity, and MoA models) to this selected subset, selecting the compounds that met the following thresholds: predicted antibacterial activity *>* 50%, the probability of toxicity *<* 0.6, and the probability of dissimilarity to known antibiotics 1 − max *p*(*c_i_*|*x*) *>* 0.5. Using these cutoffs, we obtained a list of 318 compounds. Finally, we applied additional filters based on physiochemical properties including LogP [31], TPSA [32], MPO [33], Muegge [34], and PAINS [35] such that it can maximally inform the biology and provide tractability for development into *in vivo* active chemical probes [36] and potential drug compounds (Table 4). The application of these drug-likeness filters narrowed the list down from 318 to 132 compounds. Of these 132 compounds, we were able to commercially purchase 61 molecules to experimentally validate predictions due to availability issues (Figure 3A).

**Table 4.** Physicochemical properties used as filters for narrowing down the selection of candidate molecules.

| Property | Range | Description |
| --- | --- | --- |
| Log D | $-1 < \log D < 4$ | Solubility and potential oral dosing |
| MPO | $3 < \text{MPO} < 6$ | Multiple combinations of drug-likeness factors |
| Muegge Filter | True | Multiple combinations of drug-likeness factors |
| PAINS | Remove compounds with PAINS substructure | Structural features that lead to promiscuous signals in experimental assays |
| TPSA | $< 120$ | Cellular uptake |

**Fig 3.**
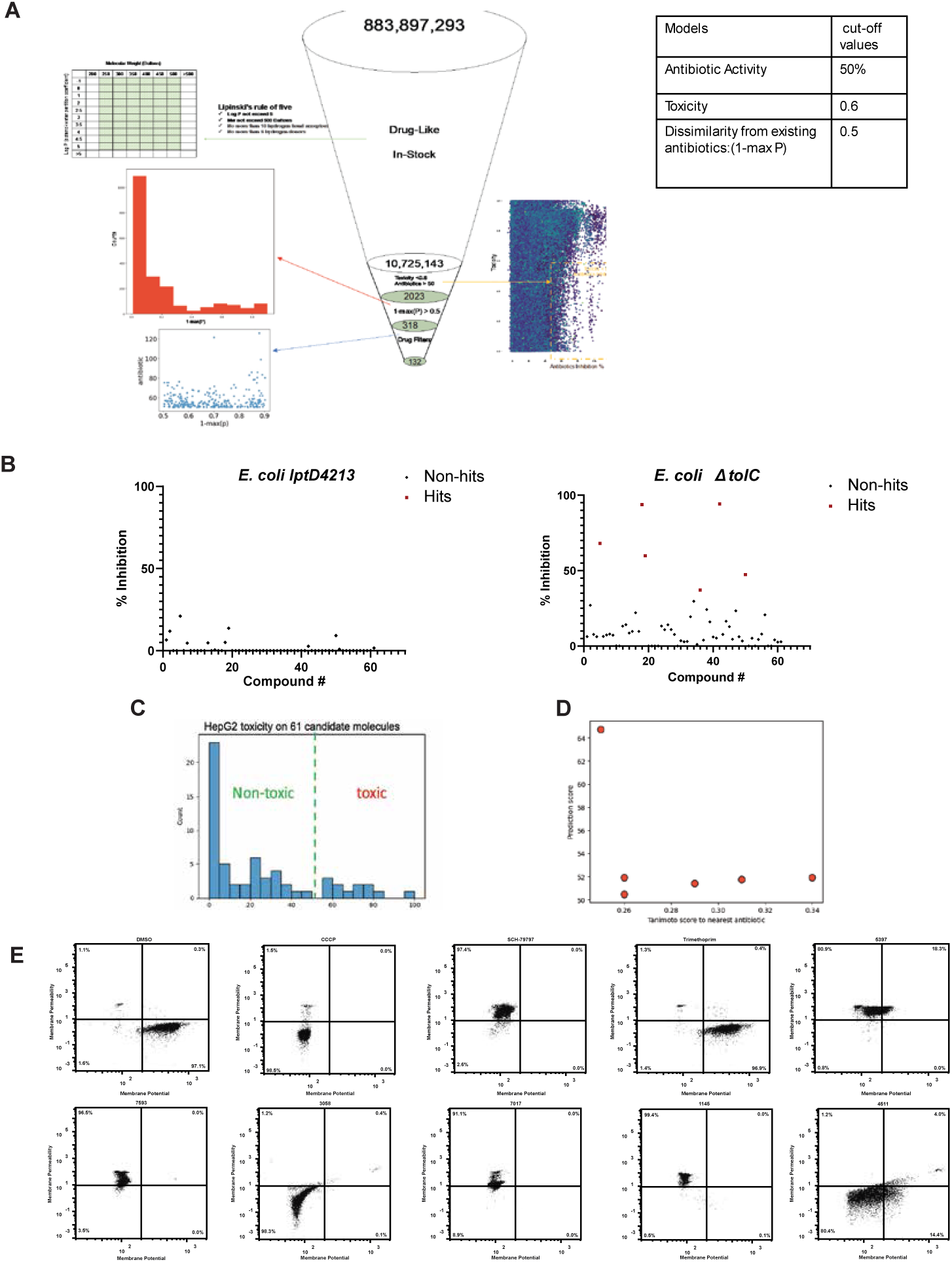
Application of our DeepChem-based model to Zinc20 database for candidate selection. (A) Workflow of the hit-to-lead strategy used for the initial *in silico* screen of Zinc20 compounds. Lipinski’s Rule of 5 was used to select the tranches from the Zinc20 listed as “in-stock.” Toxicity, antibiotic activity, and novelty thresholds were used to narrow down lead compounds. Drug-like filters were applied to generate a final list of lead compounds to validate empirically. (B) Growth inhibition calculated using optical density at 600nm (OD600)against *E. coli* lptD4213 and *E. coli* ΔtolC. Molecules with mild activity (>35%) are considered hits and labeled in red, compounds with negative inhibition values are plotted at 0. (C) Histogram of percent viability of 61 compounds against HepG2 cells after 24 hours treatment compared to vehicle-only controls. (D) Tanimoto similarity scores for 61 hits. (E) Flow cytometry of active compounds against *E. coli* lptD4213 and stained with TO-PRO-3 and DiOC2(3). Controls for vehicle only (DMSO), CCCP (depolarization), SCH-79797 (permeabilization), and not membrane active (TMP) are also included.

### Experimental validation confirms the identification of non-toxic antibacterial compounds

To validate the antibiotic activity of the initial hits, we tested their activity against *E. coli* lptD4213 and *E. coli* ΔtolC. Of the 61 compounds tested, six were able to partially inhibit the bacterial growth of at least one of the species (Figure 3B). We also tested the toxicity of these compounds against HepG2 cells at 50*µ*M. We determined that 49/61 (80.3%) of the candidate molecules were nontoxic (toxicity *<* 0.5; Figure 3C). and only one of the six compounds with strong antibacterial activity was toxic. These results demonstrate that our toxicity model successfully enriched for compounds with minimal mammalian cell toxicity. To determine if these hits represent novel chemical scaffolds, we examined the structural novelty of the six active molecules by comparing their structures to those of known antibiotics. We calculated the Tanimoto similarity of each compound’s Morgan fingerprint to the known antibiotics used to train the XGBoost classifier. We found that these molecules had low Tanimoto similarity to all known antibiotics in this dataset, and structurally did not resemble their closest Tanimoto neighbors (Figure 3D), suggesting that these could represent novel antibiotic structures.

To further assess the model and expand our hit set, we relaxed our thresholds to include compounds with a prediction of antibacterial activity *>* 40%, a probability of toxicity *<* 0.7, and a probability of dissimilarity from known antibiotics 1 − max *p*(*c_i_*|*x*) *>* 0.2. We then purchased an additional set of around 1,000 compounds, which we will refer to as the *expanded list*. From this expanded list, we obtained 15 additional hits, 9 of which inhibited the growth of *E. coli* ΔtolC by more than 80%. The remaining six hits inhibited more than 50% of the bacterial growth (Figure 4A). We found that none of 15 were toxic (Figure 4B). We also computed the Tanimoto similarity of these compounds to known antibiotics in the training dataset, and found that the hits from the expanded list have higher Tanimoto compared to the initial six hits with more stringent thresholds, though they still represent structurally distinct scaffolds. For instance, one of the hits structurally resembled nalidixic acid (Tanimoto score 0.78). This observation aligns with our model, as lowering the threshold for uncertainty in the MoA classifier led to greater similarity to known antibiotics (Figure 4C).

**Fig 4.**
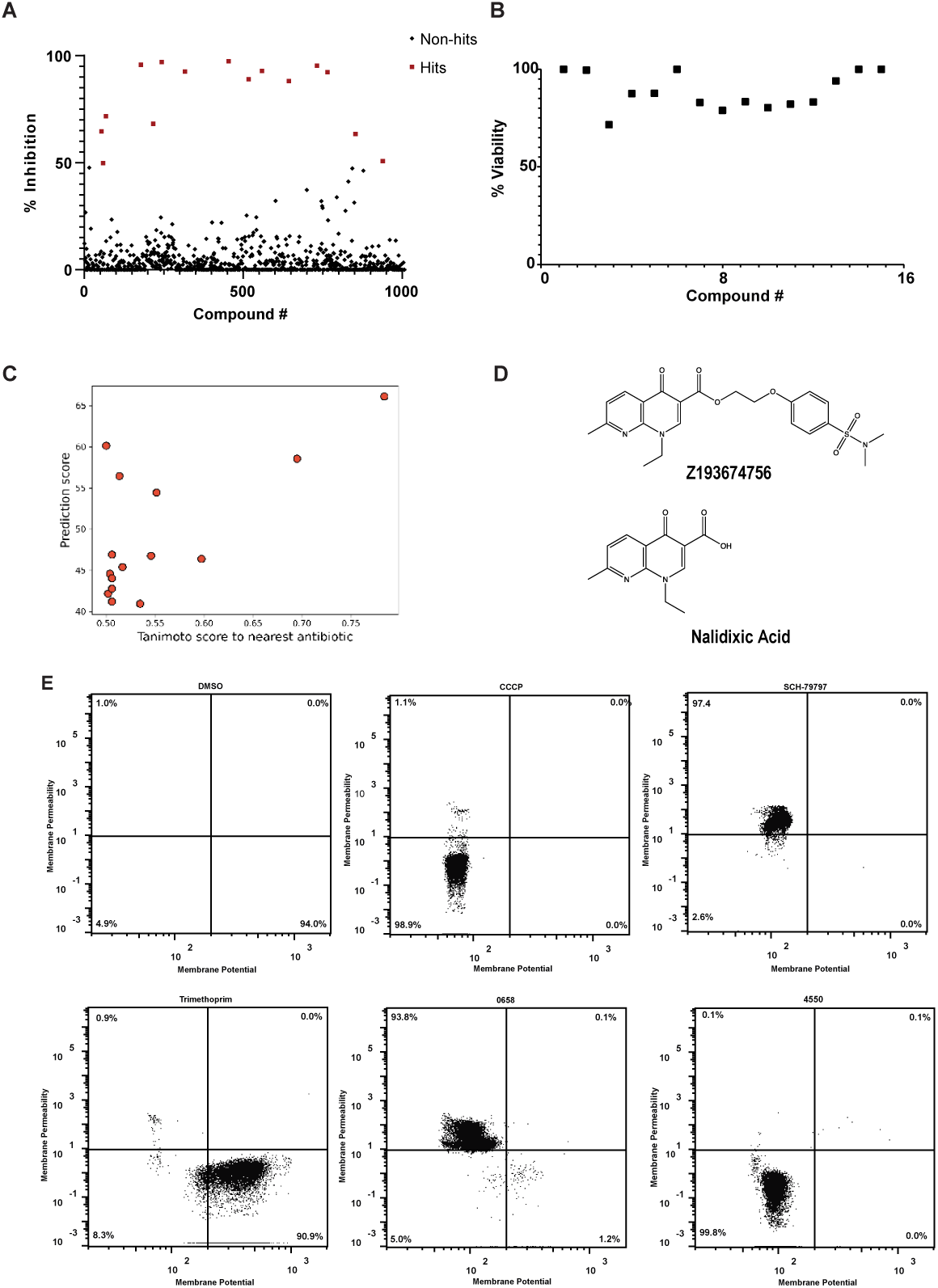
Expanded thresholds validation and proof of concept. (A) Results of the first method on the expanded list with the thresholds antibiotic activity *>* 40%, a probability of toxicity *<* 0.7, and a probability of dissimilarity from known antibiotics 1 − max *p*(*c_i_*|*x*) *>* 0.2. (B) Experimental validation of compounds for toxicity and antibiotic activity. (C) Common structural motifs present amongst hits, highlighting hit that resembles nalidixic acid. (D) Resistance mutant to confirm the target of nalidixic acid mimicking hit. (E) Flow cytometry of active compounds against *E. coli* lptD4213 and stained with TO-PRO-3 and DiOC2(3). Controls for vehicle only (DMSO), CCCP (depolarization), SCH-79797 (permeabilization), and not membrane active (TMP) are also included.

Across both screens, our pipeline successfully identified 21 compounds with antibacterial activity, the vast majority of which exhibited minimal mammalian cell toxicity and represented chemically distinct scaffolds by Tanimoto similarity metrics. These results demonstrate that our approach effectively enriches for non-toxic antibacterial compounds that are structurally dissimilar from known antibiotics.

### Identified compounds exhibit membrane-targeting activity with potential for bacterial selectivity

To determine the mechanism of action of our hits, we first performed flow cytometry on *E. coli* lptD4213 treated at 500*µ*M of each compound and stained with TO-PRO-3 and DiOC2(3), which report membrane integrity and polarization, respectively. We also included controls for vehicle only (DMSO), CCCP (depolarization), SCH-79797 (permeabilization), and non-membrane active (TMP) compounds in this assay. We found that four of the six initial hits disrupted membrane integrity while two caused membrane depolarization 3E). For the expanded list, we performed flow cytometry against E. coli lptD4213 and found that 8/15 compounds from the expanded list were membrane-active, including 2 permeabilizers and 6 depolarizers (4E).

Membrane disruption and depolarization were not MoA categories within the XGBoost model. However, the prevalence of membrane-active compounds among our hits is noteworthy for several reasons. First, these compounds exhibited selectivity between bacterial and mammalian cells as they were active against bacteria at 50-500*µ*M while showing minimal toxicity to mammalian HepG2 cells at 50*µ*M. This bacterial selectivity suggests that these molecules may preferentially target bacterial membrane compositions or architectures over mammalian membranes, a property that could be valuable for antibiotic development. Second, membrane-active antibiotics have historically been important therapeutic agents. These include daptomycin for serious Gram-positive infections [37] and polymyxins (such as colistin) as last-resort treatments for multidrug-resistant Gram-negative bacteria [38]. Identifying new scaffolds with this activity could provide starting points for developing compounds that overcome resistance to existing membrane-targeting agents. Third, the fact that our pipeline enriched for membrane-active compounds despite not explicitly training for this MoA provides important insights into biases present in our training data, as we discuss further below.

To further characterize the mechanism of action for compounds from the expanded list that were not membrane-active, we generated resistant mutants against selected compounds by plating *E. coli* ΔtolC on plates containing 4X MIC of the antibiotic. For compound Z193674756, we identified resistant mutants in *gyrA* and *gyrB*, which encode DNA gyrase (Figure 4D). This molecule contains a heterocycle with a bicyclic core motif indicative of a quinolone antibiotic similar to nalidixic acid, which is known to inhibit DNA gyrase. This finding was consistent with the higher Tanimoto similarity (0.78) observed for this compound and demonstrates that when our MoA dissimilarity threshold is lowered, the pipeline can identify derivatives of known antibiotic classes.

### Refined models with Pareto optimization or filter relaxation did not significantly improve results

Using insights learned from the first experimental iteration, we refined our models and methodology to enhance the accuracy and robustness of our machine-learning pipeline. For the antibacterial activity model, we expanded the training data to include data from our first iteration of experimentation as well as recent literature [17]. In total, we collected 124,888 data points with inhibition percentage labels and 142,815 data points with binary antibacterial activity labels (Table 5). We trained regression and classification models on these datasets using D-MPNN models [17] implemented within the Chemprop package [15]. To further improve performance, we incorporated 200 molecule-level features computed using RDKit and applied model ensembling. This resulted in notable improvement in accuracy for both activity and toxicity models. For the antibacterial activity model, the best-performing architecture was an ensemble of 20 classification models developed using Chemprop, achieving an ROC AUC of 0.75 and a precision-recall AUC of 0.17. For the toxicity model, an ensemble of five classification models developed using Chemprop emerged as the best-performing configuration, achieving an ROC AUC of 0.95 and a precision-recall AUC of 0.52.

**Table 5.** Data summary for continuous and binary data in the second method.

|  | Training/Validation | # Datapoints | % Active Compounds |
| --- | --- | --- | --- |
| Continuous Data | Training | 99910 | 0.33 |
|  | Validation | 24978 | 0.33 |
| Binary Data | Training | 114252 | 4.56 |
|  | Validation | 28563 | 4.57 |

Because we identified many membrane-active molecules from the first round of experimentation, we modified the XGBoost model by adding a new class of antibiotic mechanism called membrane active/ionophore (Table 6). We also addressed data imbalance in the XGBoost training data by creating a large ensemble with varying data-balancing ratios, hyperparameters, and initial conditions. After the changes to data and models, the new XGBoost model had an average precision of 0.96, recall of 0.95, and F1 score of 0.95, indicating that the classifier improved substantially with this additional category (Figure 5B).

**Table 6.**
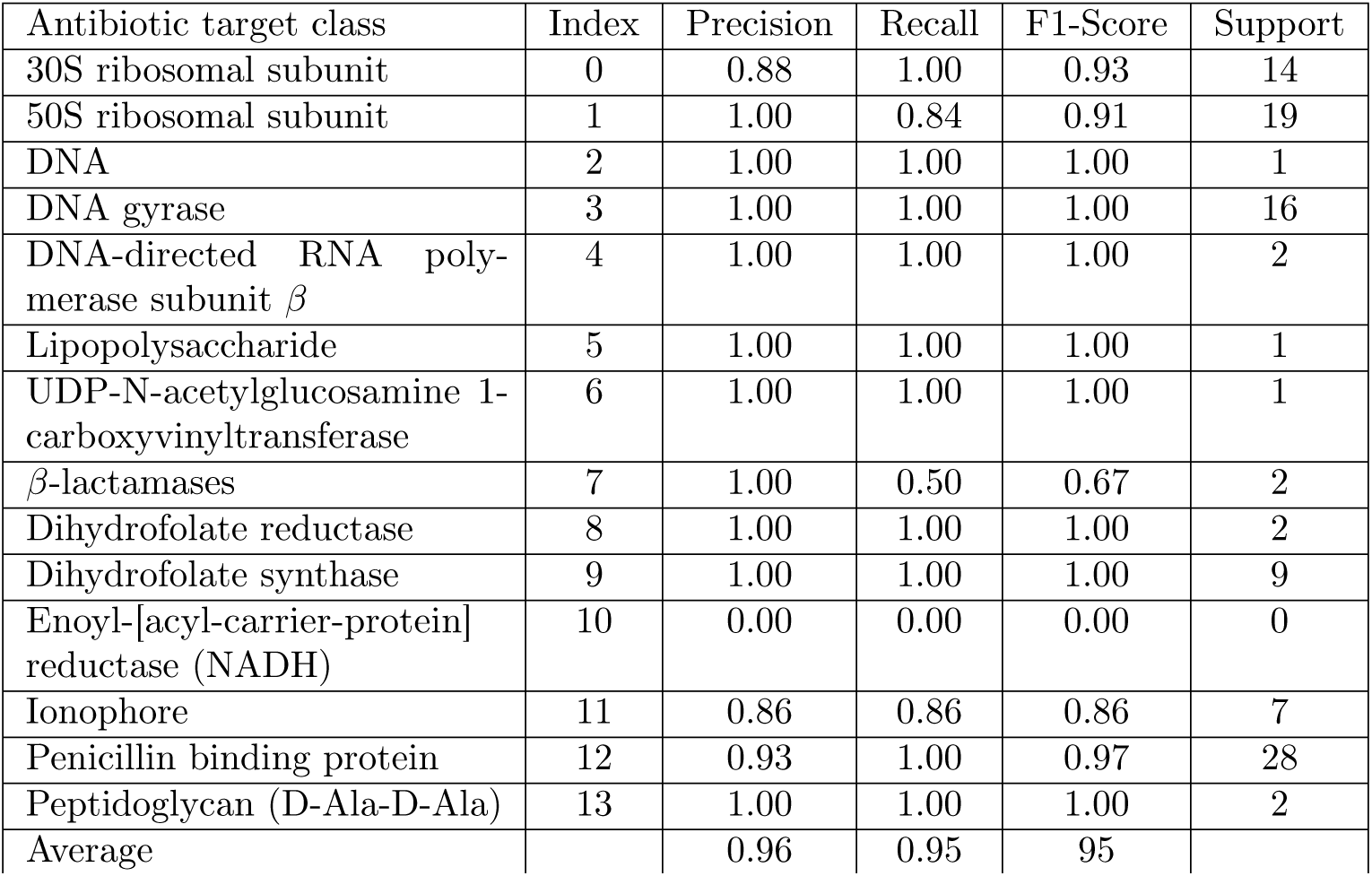
XGBoost ensemble performance on known antibiotic classes for the second approach. Individual values are given for known antibiotic categories curated from the KEGG database.

**Fig 5.**
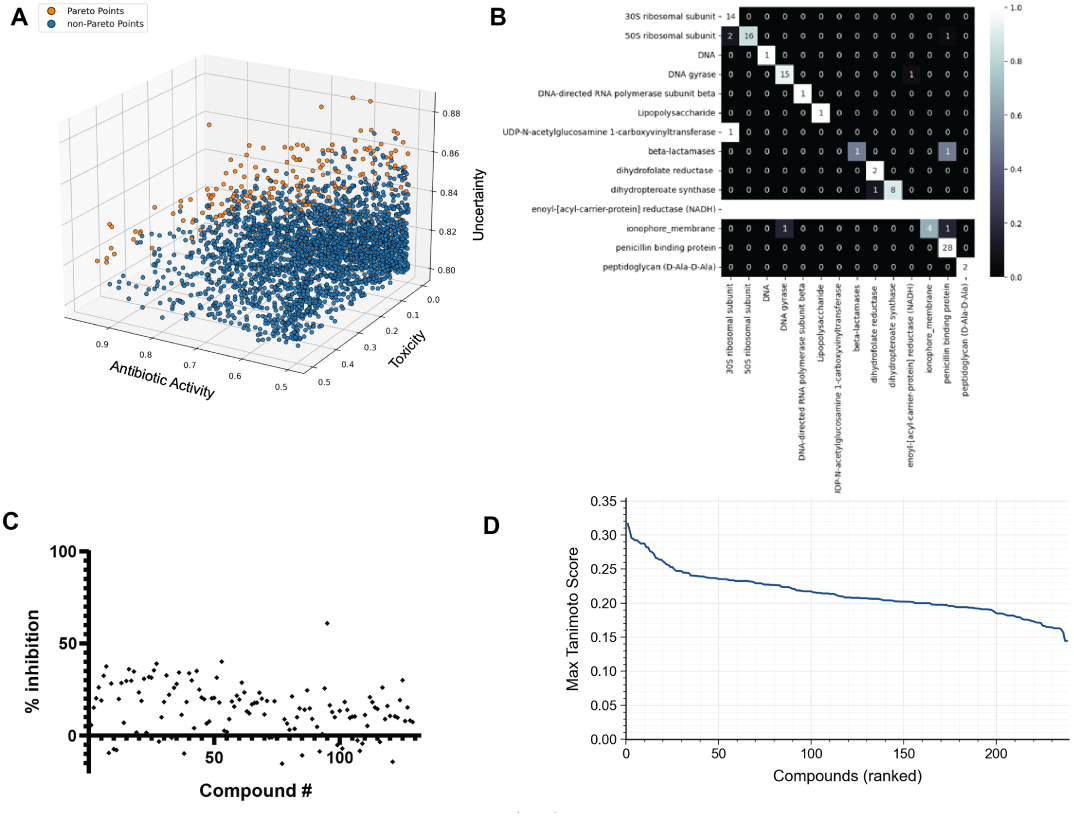
Pareto front optimization. (A) Schematic showing multi-objective optimization.(B) Confusion matrix for XGBoost model with the addition of the ionophore/membrane class of antibiotics. (C) Results of this approach. None of the identified compounds inhibit bacterial growth. (D) Max Tanimoto similarity scores of 238 Pareto-optimal compounds, ranked in descending order. All compounds scored below 0.35 against a reference set of known antibiotics, indicating structural novelty relative to existing antibiotic scaffolds.

Our goal was to identify molecules that balance three competing objectives: high antibacterial activity, low toxicity, and novel mechanisms of action. However, no single compound is optimal for all three objectives. To address this, we used Pareto front optimization to select compounds for all three objectives simultaneously rather than the top-down threshold approach used in the first method. Because Pareto fronts can include extreme trade-off points (for example, compounds with very high antibacterial activity but also very high toxicity at the edges of the frontier), we filtered such extreme values before selecting candidates. We therefore established thresholds of ≥ 50% for antibacterial activity, ≤ 0.5 for the probability of toxicity, and ≥ 0.8 for MoA uncertainty based on the distribution of uncertainty scores for hits in the training data (Extended Data Figure 2). Applying these three thresholds to the Zinc20 database, we identified approximately 3,000 candidate compounds. To determine the top candidates, we used non-dominated sorting implemented in the pymoo package [39], which resulted in 238 Pareto optimal compounds (Figure 5A). These compounds looked promising computationally with near-zero toxicity prediction probabilities, high activity prediction scores, and all MoA uncertainty probabilities above 0.8. These compounds also looked structurally different from any known antibiotics in the training dataset by Tanimoto similarity (Figure 5D).

However, when tested experimentally, only one of our Pareto-optimized compounds showed antibacterial activity greater than 50% against *E. coli* Δ tolC (Figure 5C). Moreover, this molecule was only weakly active at the highest concentrations tested. This disappointing result, despite improved models and more sophisticated optimization methods, further reinforces our understanding that the fundamental limitation lies not in the model architecture or selection strategy, but in the mechanistic composition of the training data itself. The activity model continued to favor compounds similar to those in the training set, making it difficult to explore truly novel chemical and mechanistic space.

We also explored the possibility that our drug-like property filters (such as following Lipinsky’s rule of five and maintaining a LogP less than 5) were preferentially discarding molecules with novel mechanisms of action. To this end, we used the same Pareto optimality thresholds as above but relaxed drug-like properties to identify an additional 1285 compounds for experimental testing. Of these, 119 (9.3%) showed significant antibacterial activity (*>*80% growth inhibition of E. coli Δ tolC and only 14 (1.1%) were toxic to HepG cells. Of these 119 candidates, 15 had an MIC of 16 *µ*M or less against E. coli Δ tolC, and these 15 compounds fell into 7 chemical subtypes (Extended Data Figure 3A,B). Flow cytometry assays revealed that all 7 of these compounds acted as membrane depolarizers (Extended Data Figure 3C). These results reinforce the conclusion that our models perform well at enriching for compounds that selectively target bacterial membranes and that the lack of identifying novel protein-targeting compounds is not due to drug-like filtering.

### Analysis of mechanism-of-action bias in training data

While our pipeline successfully identified non-toxic antibacterial compounds with novel chemical scaffolds, the predominance of membrane-active compounds among our hits raised an important question: why did we not identify compounds with novel protein targets? To address this, we performed a systematic analysis of mechanism-of-action representation in our training data, and the distribution of known antibiotic MoAs in both our training data and screening hits using the (retrained) XGboost.

A subset of the active compounds in our training data are similar to known antibiotics. However, the various compounds do not equally capture the different MoA classes. This imbalance, combined with the modest recall of our model, may result in a bias toward the overrepresented MoA classes. We examine this potential bias systematically by applying the MoA model to a set of active compounds with a known MoA. We find a strong imbalance between MoA classes (6a). For example, while our model successfully learned to predict quinolones, it completely fails to predict compounds belonging to smaller classes (e.g., DHFR inhibitors) and exhibits lower than average recall for larger MoA classes such as beta-lactams, aminoglycosides, and tetracyclines. In Figure 6b. We show the distribution of XGBoost classifications for compounds with *P_max_ >* 0.5 for our training actives. A large subset of the compounds is predicted to be membrane-active, another significant portion is similar to known compounds of various classes, and a 3rd portion is not well classified by XGBoost.

**Fig 6.**
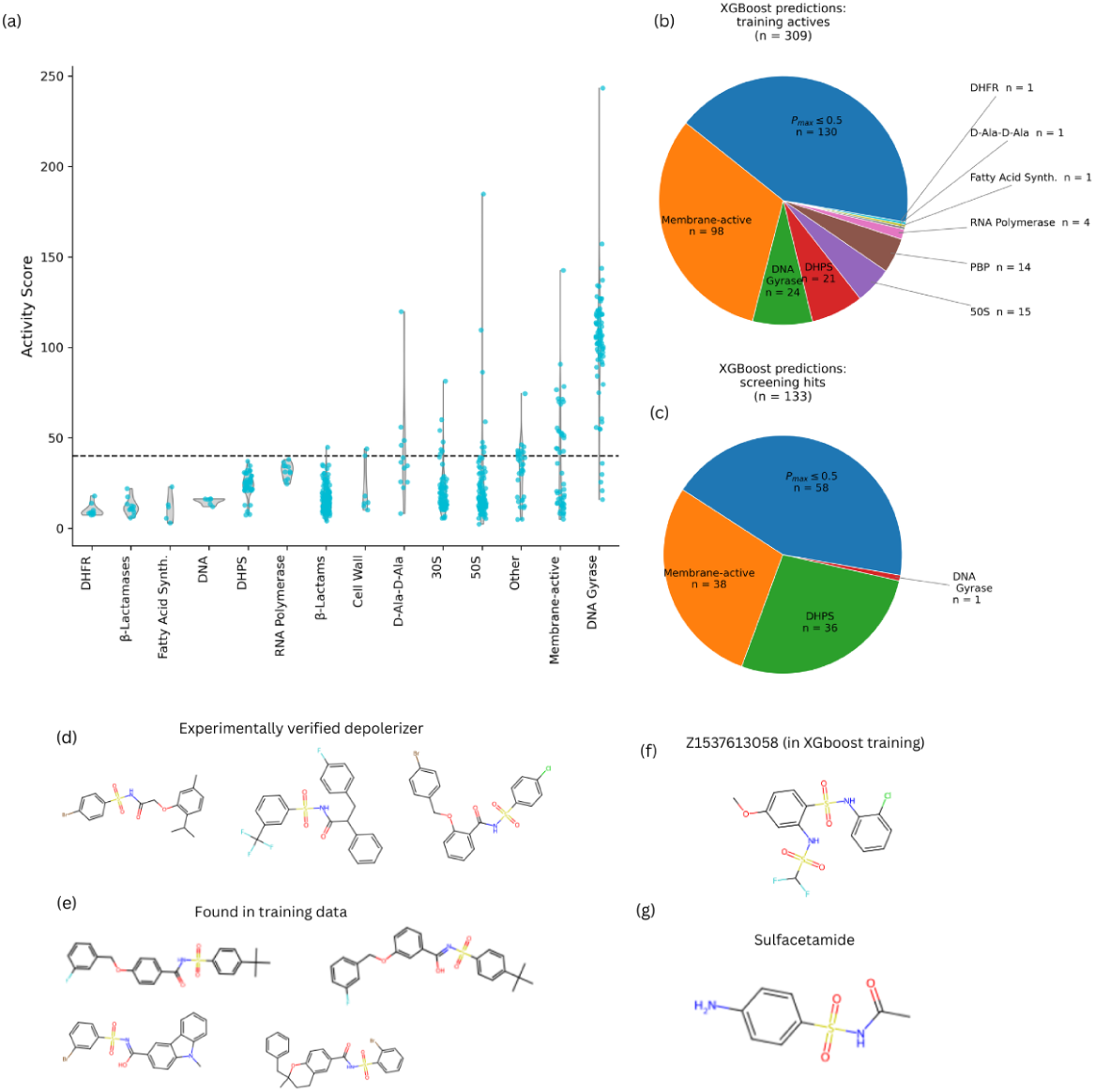
Mechanism of action performance. (A) Activity score for known compounds with a labeled MoA. The dashed horizontal line represents the expanded activity threshold. Only some of the MoA classes have any compounds that above the prediction threshold. (B) XGBoost class distribution for 309 actives in the training data. (C) XGboost class distribution for 133 actives from the our expanded screen. (D) Experimentally verified depolarizers from the screen. These have a sulfanimide-like structural moiety. Such compounds are typically classified as either membrane active or DHPS inhibiting by XGBoost (E) similar compounds found in the training data. (F) a membrane active compound added to the training data of XGBoost after the first round. (G) A known DHPS inhibitor.

Moreover, compounds classified as DHPS inhibitors and membrane-active are over-represented compared to the training data. When analyzing the structures of our novel hits, the reason for this becomes clear. In Figure 6d and 6e, we show experimentally verified depolarizer hits from our screen, and compounds found in our training data, respectively. These share a common moiety with both Z1537613058 (Figure 6f), one of the 1st round membrane active compounds added to XGboost, and with known DHPS inhibitors (e.g., Sulfacetamide; Figure 6g). This structural similarity results in confusion between these 2 classes as the top classifications. Indeed, a large majority of the screening hits (expanded threshold and drug filter removed) had either membrane-active or DHPS as their top 2 classifications. We examined whether this structural similarity resulted from a true mechanistic origin by growing *E. coli* in the presence of these compounds with TMI supplementation, but found no significant shift in the MIC. We conclude that all compounds sharing this structural moiety in the training data are likely to share the same membrane-active mechanism as our experimentally verified depolarizers. This MoA bias in the training data provides a mechanistic explanation for our experimental observations. The antibacterial activity model learned to assign high activity scores predominantly to molecules similar to those encountered during training. Since active molecules in the training data often fall within the scope of known mechanisms of action and membrane-active compounds appear to be overrepresented in antibacterial activity screen data, any molecule with a potentially novel protein target likely received a low activity score and was likely discarded from our search.

## Discussion

In this work, we developed a machine learning pipeline that addresses chemical novelty in antibiotic discovery by using mechanism-of-action classification upfront rather than as a post-hoc filter. Applied to the Zinc20 database, our approach successfully identified non-toxic antibacterial compounds that are structurally distinct from known antibiotics by Tanimoto similarity metrics. Of the initial 61 compounds tested, 6 showed antibacterial activity, with only 1 displaying toxicity against mammalian cells. By relaxing our thresholds, we identified 15 additional hits with strong growth inhibition activity. These results demonstrate that our pipeline effectively enriches for compounds with favorable therapeutic properties, namely antibacterial activity coupled with minimal mammalian toxicity.

A notable finding from our work is that the majority of identified hits exhibited membrane-targeting activity. While membrane disruption and depolarization were not explicit categories in our original XGBoost MoA classifier, these compounds displayed selectivity between bacterial and mammalian cells, remaining active against bacteria while showing minimal toxicity to mammalian cells at the same concentrations. This bacterial selectivity is particularly significant given the historical importance of membrane-active antibiotics such as daptomycin and polymyxins [37, 38]. The identification of new chemical scaffolds with membrane-targeting activity and demonstrated bacterial selectivity could provide valuable starting points for developing next-generation membrane-active antibiotics that overcome resistance to existing agents. This is relevant given that bacteria-selective membrane-active compounds represent a largely underexplored approach in modern antibiotic development.

Despite these promising findings, our pipeline did not achieve its primary goal of identifying compounds with novel protein targets, even with model refinement and drug-like filter relaxation. Our systematic analysis revealed that this limitation stems both from representation biases in the training data and from MoA biases in the antibacterial activity model. The activity model learned to assign high activity scores predominantly to molecules similar to specific classes in the training dataset, which is enriched for certain mechanistic types, specifically membrane-active compounds. This creates a contradiction: while our XGBoost MoA classifier was designed to prioritize mechanistically novel compounds, the activity model simultaneously deprioritized these same compounds because they differed from patterns learned during training. This observation highlights critical challenges for structure-based machine learning approaches in antibiotic discovery: predictive models can only effectively explore the chemical and mechanistic space represented in their training data, and they are likely to significantly reduce the mechanistic diversity compared to the original training data.

Our second-round efforts, which incorporated improved model architectures (D-MPNN ensembles), expanded training datasets, additional molecule-level features, an enhanced XGBoost model with a membrane-active category, and Pareto optimization, yielded only one weakly-active hit from 238 computationally promising candidates. This result reinforces the idea that the fundamental limitation is not computational but empirical, and specifically stems from the scarcity of mechanistically diverse, well-annotated training data. The field currently lacks comprehensive datasets of antibacterial compounds with validated mechanism-of-action labels spanning diverse protein targets and pathways. It is also possible that adding compounds with partial antibacterial properties but diverse MoAs to the training data will allow ML models to traverse the mechanistic search space without relying on effective antibiotics at every point of the space; similarly, toxic compounds could also be used to train the MoA models. Without comprehensive training data, machine learning models, regardless of architecture, will continue to recapitulate patterns from existing chemical classes of successful antibiotics rather than genuinely explore novel mechanistic space.

Our findings suggest several directions for advancing machine learning-based antibiotic discovery. First and most critically, the field needs substantial investment in generating mechanistically annotated training datasets. This requires systematic experimental characterization of compound mechanisms across diverse chemical scaffolds, including compounds that target underrepresented pathways and proteins, separately from their antibacterial and toxicity profiles. Such datasets would enable activity models to learn representations of antibacterial activity that extend beyond currently prevalent mechanisms.

Second, alternative approaches to measuring chemical novelty beyond Tanimoto similarity merit exploration. Metrics such as the Dice coefficient [40], which is more sensitive to shared features, the maximum common substructure (MCS) similarity [41], which effectively identifies shared scaffolds despite computational intensity, and the Tversky index [42], which provides flexible weighting of shared and unique features, could help reduce mechanistic redundancy in candidate selection. Additionally, explicit filters for common structural motifs such as quinolone cores could prevent selection of obvious derivatives.

Third, alternative molecular featurization approaches could potentially improve exploration of chemical space. Models that combine multi-layer neural network embeddings of molecular fingerprints with GCN embeddings [43] could capture complementary features, while CNNs using image representations of molecules [44] may provide novel perspectives on molecular structures that help models escape overreliance on training data patterns.

Finally, our observation that the activity model preferentially assigns high scores to molecules resembling training data suggests an alternative strategy: rather than attempting to simultaneously optimize for activity, toxicity, and novelty, future approaches might prioritize identifying novel, non-toxic scaffolds first, then use medicinal chemistry to optimize antibacterial activity and drug-like properties. This inverted workflow could better balance the competing objectives and necessary exploration inherent in antibiotic discovery.

While our pipeline did not achieve its primary goal of discovering antibiotics with novel protein targets, it demonstrates clear utility for related applications. The approach could effectively identify derivatives of known antibiotic classes with enhanced potency or specificity against particular pathogens. It could also be adapted for optimization of inhibitors targeting specific enzymes or pathways, where training data can be more readily generated for a single target. For molecular optimization (improving existing compounds rather than discovering entirely novel scaffolds) this framework provides a robust foundation.

An additional consideration is that our pipeline used Lipinski’s Rule of Five to filter for drug-like molecules. While these rules predict oral bioavailability in humans [30], they may not accurately predict bacterial uptake and accumulation, particularly given the barrier posed by the Gram-negative outer membrane. Alternative frameworks, such as the eNTRy rules [45] or accumulation rules [46] developed specifically for antibacterial compounds, may enable exploration of different chemical spaces more appropriate for bacterial targets. However, relaxing these physicochemical constraints would require careful consideration of the balance between expanding accessible chemical space and maintaining eventual drug development tractability.

In conclusion, our work demonstrates both the promise and current limitations of machine learning for antibiotic discovery. We successfully developed a pipeline that enriches for non-toxic, active compounds, including membrane-targeting agents with demonstrated bacterial selectivity. However, the identification of compounds with truly novel protein targets remains elusive, constrained not by computational methodology but by the mechanistic diversity of available training data. Advancing the field will require concerted effort to generate comprehensive, mechanistically annotated datasets spanning diverse chemical and biological space. Only with such foundational resources can machine learning fulfill its potential to accelerate the discovery of the next generation of antibiotics.

## Methods

### Data Processing

#### First Method: Data processing for antibiotic activity regression model

To prepare for the training data for the antibiotics and toxicity GCN models, we first converted the original SMILES expression to International Chemical Identifier (InChI) using the RDkit [47]. The unique InChI expressions were used to remove the duplicates within the raw training data and the duplicates in the testing data. Compounds consisting of rare atoms (As, Se, Ca, Na, Li, Fe, Cu, Hg, Pt, Zn, Au, Ac, Bi, Ge, Te, Co, Re, Pb) were further removed for model stability. Finally, RDkit was used to convert InChI expression back to the SMILE expression as the input strings for better consistency. After data cleaning, a total of 122,851 compounds were used as the training and validation data for the antibiotics GCN model. To evaluate this model, we used a library containing 2,208 compounds with binary antibiotic scores tested in-house from a subset of the Chembridge DIVERSet Library as the test data.

#### First Method: Data processing for toxicity classification model

After data cleaning, a total of 50,707 compounds were used as the training and validation data for the toxicity GCN model. To evaluate this model, we used a set containing 385 compounds with binary toxicity scores as the test data.

#### Second Method: Data processing for antibiotics regression model

To develop the regression model, it is essential that the training data consist of continuous values. For this purpose, we integrated compounds containing continuous inhibition rates obtained through experimental screenings from three distinct sources. The first source includes the inhibition rates measured from in-house screenings conducted in rich media. The second source is the publicly available Shared Platform for Antibiotic Research and Knowledge (SPARK) and COADD databases [25]. The third source comprises the inhibition rates of additional compounds incorporated after an initial round of predictions. After data pre-processing any duplicate compounds had their experimental values averaged if the range of these values was less than 50%. After deduplication, the initial screening dataset contained 51,995 compounds, the publicly available dataset contained 76,182 compounds, and the new screening dataset contained 1,070 compounds. This resulted in a final training dataset of 124,888 compounds.

#### Second Method: Data processing for antibiotics classification model

To develop the classification model, we combined data from multiple sources containing both binary and continuous values. The first source includes inhibition rates measured from in-house screenings. These values were binarized using an 80% threshold. For compounds with only OD600 values, a threshold of 0.5 was used for binarization. Secondly, compounds from publicly available databases were included with binarization thresholds of 80% inhibition rate and less than 30 *µ*g/mL MIC, respectively. Additionally, all compounds in the Kyoto Encyclopedia of Genes and Genomes and classified as antibiotics from other manuscripts [17]were included as positive compounds. We also included inhibition rates of additional compounds incorporated after an initial round of predictions, binarized with an 80% threshold. After data pre-processing, in each dataset, duplicate compounds were removed if they had both positive and negative values. After deduplication, the initial screening dataset contained 52,062 compounds. The data publicly available contained 76,182 compounds with inhibition data and MIC values for 15,976 compounds, respectively. The Kyoto Encyclopedia of Genes and data from other publications contained 7510 compounds. The new screening dataset contained 1,070 compounds. This resulted in a final training dataset of 142,790 compounds.

### Prediction methods for antibiotic screens

#### First Method: Prediction method for antibiotic activity and eukaryotic toxicity

Two separate graph convolutional network (GCN) models were trained using the DeepChem toolkit to independently predict the antibacterial activity and toxicity of compounds based on their molecular structures. The two graph neural networks were used to learn to predict the toxicity and antibacterial activity of molecules. The GCN models were trained on data with the molecular structure of small molecules represented by the SMILES strings. This modeling approach focuses on transforming the SMILES representation into a graph representation where each node is an atom defined by its fingerprints and the bonds are edges of the graph. This modeling strategy primarily focuses on atom features as the main representation of the molecule and executes the message-passing operation through neighborhood aggregation over the nodes of the graph (atoms). To train the model to predict antibacterial activity in the regression mode, the training data were first splitted into the training set and validation set in the percentage of (80, 20), and each split has a proportional number of the active compounds. The compounds in these datasets were defined as active if their inhibition percentage exceeded the 80% threshold. Since the training dataset was imbalanced, weights were added to balance the numbers of active and inactive compounds. To train the models, the hyperparameters including batch size, the structure of the GC layers and the dense layers, dropout rate and learning rate, were determined using the “hyperopt” python library with the default algorithm. Then the GCN for antibacterial activity prediction was built with two graph convolution (GC) layers both with 128 nodes and a dense layer with 64 nodes. The batch size, dropout rate and learning rate were selected as 100, 0.205, and 0.0007. On the other hand, to train the GCN model to predict toxicity in the classification mode, the training data were also split into the training and validation set in the percentage of (0.8, 0.2) with each split containing the same percentage of active compounds. Sample weights were also added again for the imbalanced data. After the hyperparameter optimization using “hyperopt”, the GCN model for toxicity prediction was constructed with two GC layers both with 128 nodes and a dense layer with 16 nodes. The batch size, dropout rate and learning rate were set as 64, 0.304, and 0.0006 individually.

#### First Method: Measuring similarity to existing antibiotics

We developed a machine learning model using the XGBoost method to predict antibiotic target classes. We used the initial dataset from KEGG [29] and performed several data preprocessing steps. These steps removed datapoints with missing targets and deleted rows associated with underrepresented targets such as “alanine racemase” and “ATP synthase subunit C”. We also merged similar classes, like “ribosome” into “50S ribosomal subunit”, “D-alanine-D-alanine ligase” into “peptidoglycan (D-Ala-D-Ala)” and “DNA gyrase topoisomerase IV” into “DNA gyrase”. These processes resulted in a clean dataset of 482 antibiotics with their corresponding targets in thirteen categories. Next, we converted the antibiotics’ SMILES representations, which obtained from the InChI identifiers, into Morgan fingerprints with a radius of 2. We also converted the antibiotic target classes to numerical values. Subsequently, we split the dataset into the training and test sets at an 80:20 ratio. Finally, we employed an ensemble of XGBoost multi-class classification models to improve prediction accuracy using Python’s Scikit-learn library. The ensemble consists of twenty fine-tuned XGBoost models, each with different initialization, yet trained on the same dataset. For predictions, the ensemble averages the probabilities across all the individual models.

#### Second Method: Prediction method for antibiotic activity and eukaryotic toxicity

In the second approach, we used a (D-MPNN) implemented in Chemprop to predict antibiotic activity and toxicity from molecular SMILES strings, which were converted into graph representation using RDKit. The model performs several message-passing steps, aggregating information from neighboring atoms and bonds to represent local chemistry. During each step, the vector feature of a bond is updated by summing the features of neighboring atoms and bonds, applying a neural network layer, adding the bond’s previous featurs, and applying ReLU activation. After a set number of steps, the updated features are summed to create a single representation for the entire molecule. This representation is subsequently processed by a feedforward neural network to predict the target properties: antibiotic activity and toxicity. The D-MPNN models differ from the GCN models used in the first method by treating the molecular graph as a directed graph and incorporating information about both atoms and bonds into the feature representation of molecules. To improve predictive performance, we enhanced the architecture through hyperparameter optimization, integrating molecule-level features and ensembling. A hyperparameter optimization grid search was performed over the following parameters: depth, feedforward number of layers, number of hidden layers, feedforward hidden size, and dropout (see Table Extended Data Table 3), for 50 epochs. To address the challenge of extracting global molecular features in large molecules, we combined the molecular representation obtained from message passing with 200 additional molecule-level features computed using RDKit. Model robustness was further enhanced through ensembling; we trained 20 independent models for antibiotic activity and five models for toxicity predictions, each initialized with different random weights but trained on identical dataset splits. Final predictions were obtained by averaging outputs from the ensemble models.

#### Second Method: Measuring similarity to existing antibiotics

In the second round, we expanded and enhanced the XGBoost dissimilarity model from different angles. After the first round of the dissimilarity model, we determined the necessity of adding a group that targets membranes or functions as ionophores. Therefore, we added new molecules for the ianophore class resulting in 14 classes. After data cleaning, we obtained a dataset of 570 molecules. Despite the increase in size, certain classes (e.g., DNA-directed RNA polymerase subunit beta) remained significantly underrepresented, while others (e.g., penicillin-binding protein) had considerably more data points (up to 1,390). To address this imbalance, we tested various up-weighting ratios for underrepresented classes relative to the most populous class. We implemented an up weighting approach to ensure each class receives at least *r*% of the weight assigned to the majority class, with *r* ∈ [0.1, 0.2, 0.3, 0.4, 0.5, 0.6, 1]. At *r* = 1, all classes are fully balanced, regardless of their individual data point counts. For each value of *r*, we trained and optimized model hyperparameters across multiple randomized initializations. Models with an F1 score above 0.94 were selected to form our final ensemble. In this selection process, we evaluated the models’ ability to differentiate between data points from known and unknown classes, regardless of correct class identification among known classes. This evaluation served as an additional filter to include models that effectively distinguish novel classes and exclude overconfident models. Our final ensemble model included 19 diverse models and achieved an F1 score of 0.95, a precision of 0.96, and a recall of 0.95.

#### Evaluations of the three models

After the three models were developed (two GCN models and an XGBoost model), we employed two different strategies to assess their performances. For the two separate GCN models predicting antibacterial activity and toxicity, we computed the area under curve (AUC) of receiver operating characteristic (ROC) Curves and Precision-Recall (PR) curves. ROC curves summarize the trade-off between true positive rate (TPR) and false positive rate (FPR), while PR curves summarize the trade-off between precision and recall. To compute TPR, FPR, precision and recall, the continuous values of the test data for the antibacterial activity model were first binarized with the threshold 80%, and the binary values of the test data for the toxicity model were directly used. To evaluate the XGBoost model for measuring the similarity to existing categories of antibiotics, we first assessed the performance of the trained XGBoost classifier to known antibiotics categories by computing the precision, recall and F1-score of the test dataset.

The two scores indicated that our classifier could successfully classify compounds based on the molecular fingerprints. However, our goal to train the XGBoost model is to prioritize compounds that carry novel mechanisms in killing bacteria (i.e. compounds that are structurally and functionally dissimilar to existing antibiotics). Therefore, we defined the dissimilarity score 1 − max *p*(*c_i_*|*x*) ; where *i* ∈ {1*, . . . ,* 14} antibiotics categories to indicate the level of dissimilarity to well-known antibiotics. To examine if the dissimilarity score we defined is a valid metric, we applied the XGBoost model on the Morgan fingerprints of 7 molecules we excluded in the training step and a molecule that was randomly generated. Higher dissimilarity scores of the 8 molecules which should be structurally and functionally dissimilar to the known categories of antibiotics supported the validity of the dissimilarity scores.

### Validation of lead compounds

#### Validation of antibiotic activity

Overnight cultures were diluted 1:200 in LB Broth and added to a 96-well plate. Small molecules were added at 50*µ*M final concentration in 0.5% DMSO to wells and grown at 37°C with continuous shaking for 16 hours. Cell growth was measured by optical density (OD600). Assays were performed in Tecan InfiniteM200 Pro (Männedorf, CH) microplate readers.

#### Validation of toxicity

HepG2 cells were seeded in collagen-coated 96 well plates at a density of 1 × 10^5^. Compounds were added to cells at 50*µ*M. and compared to HepG2 cells treated with 0.5% DMSO. After 24 hours, cell viability was assessed using Alamar Blue HS following manufacturer instructions, measuring the emission of fluorescence at 590nm. To calculate the percentage of toxicity we used the following formula:

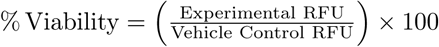

#### Flow cytometry

Overnight *E. coli* lptD4213 cultures were diluted 1:100 and grown to early-mid exponential phase (OD600 = 0.4-0.6) at 37°C. Each culture was then diluted 1:10 into PBS and treated with the desired concentration of antibiotic for 15 minutes. Cells were then stained with the BacLight Bacterial Membrane Potential kit (ThermoFisher B34950). This kit uses DiOC2(3) to measure a cell’s membrane potential as a ratio of green (488 nm ex, 525/50 nm em) to red (488 nm ex, 610/20 nm em) [48]. Membrane integrity was measured by staining cells with TO-PRO-3, a dye that is excluded from cells with an intact membrane (640 nm ex, 670/30 nm em). The LSRII flow cytometer (BD Biosciences) at the Flow Cytometry Resource Facility, Princeton University, was used to measure the fluorescent intensities of both dyes in response to antibiotic treatment. 100,000 events were recorded for each data file. Data was analyzed using FlowJo v10 software (FlowJo LLC, Ashland, OR).

#### Resistance mutation identification

To isolate resistant mutants 10^8^ colony forming units of *E. coli* ΔtolC. were plated on LB agar plates containing 4× the minimum inhibitory concentration (MIC) of drug. MIC was determined as the minimum concentration at which no bacterial growth was observed over 16 hours at 37°C. Plates were grown at 37°C for 48 hours, after which individual colonies were picked and grown at 4× MIC of antibiotic in LB broth to confirm resistance.

To identify mutations able to confer resistance, whole-genome sequencing was performed and compared to the parental strain of *E. coli* ΔtolC. Genomic DNA was extracted using the DNeasy blood and tissue kit (Qiagen 69504) and sequenced using an Illumina NextSeq 2000 system at SeqCenter.

### Tanimoto of Morgan Fingerprint calculations

RDKit [47] was used to generate the Morgan fingerprints for each small molecule using a radius of 2 and 2048-bit fingerprint vectors. These Morgan fingerprints determine all the substructures centered around an atom with a radius size 2 away from the central atom. The Tanimoto similarity is the total number of shared substructures between two molecules divided by the total number of substructures. A higher Tanimoto similarity (closer to 1) indicates that two molecules are more alike, where a score of 0 would indicate that no substructures are shared.

## Acknowledgments

We would like to thank Matthew Cahn, Senior Research Systems Analyst in the Biology Department at Princeton University, for his assistance with running computational jobs on the Della cluster in the CryoEM partition at Princeton University. This work was supported in part by the Zuckerman STEM Leadership Program and by Yad Hanadiv Foundation. S.B., C-Y.W. and B.E. were funded in part by grants from the Parker Institute for Cancer Immunology (PICI), the Chan-Zuckerberg Institute (CZI), the Biswas Family Foundation, NIH NHGRI R01 HG012967, and NIH NHGRI R01 HG013736. B.E. is a CIFAR Fellow in the Multiscale Human Program.

B.E. is on the Scientific Advisory Board for ArrePath, Inc., GSK AI for Cancer, and Freenome. Z.G. is the Founder of ArrePath, Inc.

## Author contributions statement

Z.G. and B.E. conceived the experiments, C.C., S.G., S.B., J.S, I.I. and C-Y.W. conducted the experiments, C.C., S.G., S.B., J.S, I.I., H.K., B.E., and Z.G. analyzed the results. All authors reviewed the manuscript.

## Extended Data

The code and data used for this study are available at https://github.com/PrincetonUniversity/Novel-Antibiotic-Development-Using-AI.

**Extended Data Table 1:** Hyperparameters of the GCNs of the first method (the DeepChem models)

| Hyperparameter | Antibiotic Model | Toxicity Model |
| --- | --- | --- |
| Number of atom features | 78 | 78 |
| Graph convolutional layers | [128, 128] | [128, 128] |
| Dense layer size | 64 | 16 |
| Dropout rate | 0.205 | 0.304 |
| Mode | regression | classification |
| Learning rate | 0.0007 | 0.0006 |
| Batch size | 100 | 64 |

**Extended Data Figure 1:**
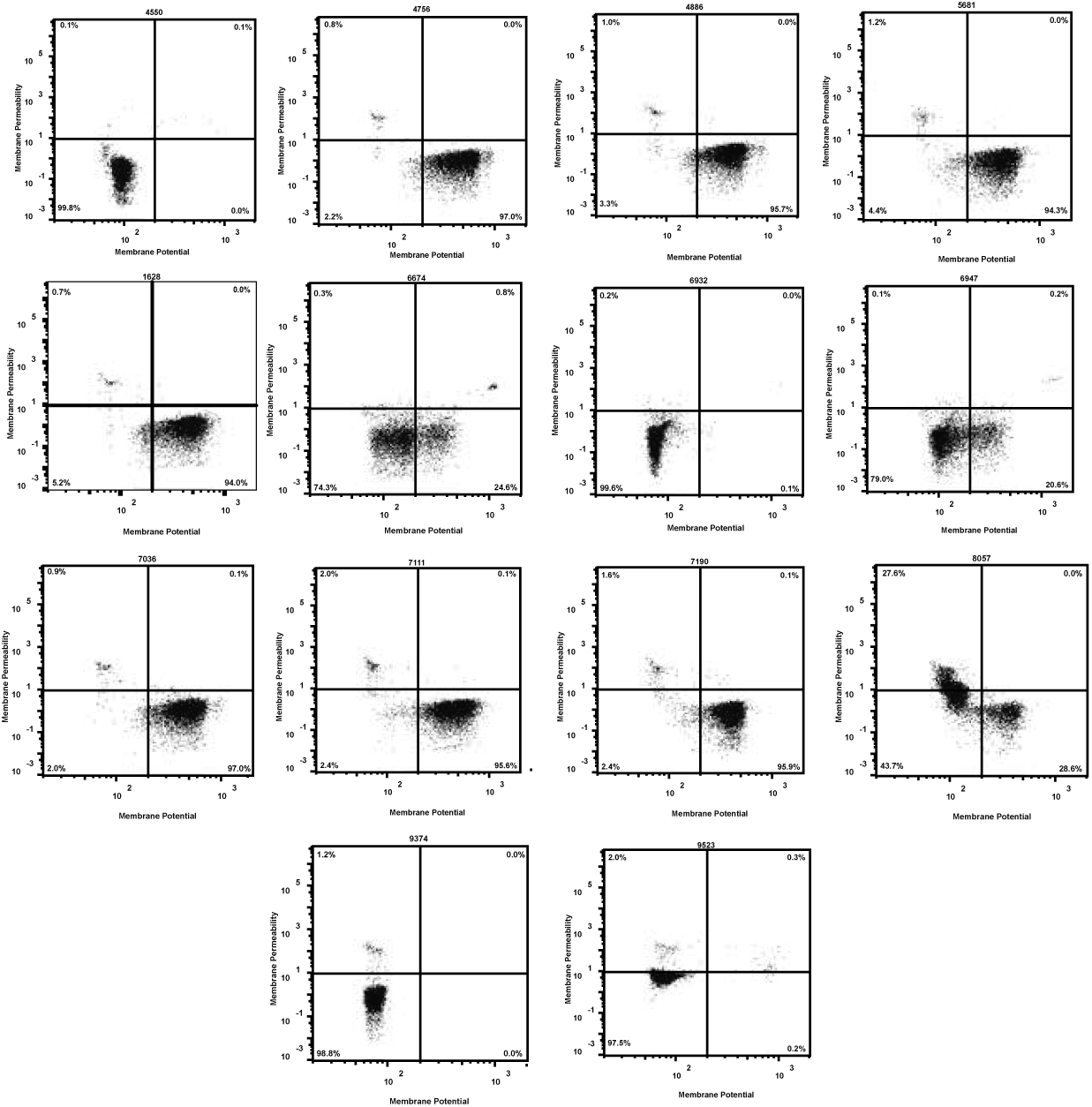
Flow cytometry of active compounds against *E. coli* lptD4213 and stained with TO-PRO-3 and DiOC2(3). Controls for vehicle only (DMSO), CCCP (depolarization), SCH-79797 (permeabilization), and not membrane active (TMP) are also included.

**Extended Data Table 2:** Hyperparameters of the GCNs of the first method (the Chemprop models).

| Parameter | Antibiotic Model | Toxicity Model |
| --- | --- | --- |
| Batch size | 20 | 15 |
| Depth | 5 | 6 |
| Dropout | 0 | 0.3 |
| ffn Hidden size | 1200 | 1700 |
| ffn num layers | 3 | 3 |
| Final learning rate | $5.40 \times 10^{-6}$ | $9.81 \times 10^{-6}$ |
| Hidden Size | 1200 | 1700 |
| Init learning rate | 0.00011 | $4.98 \times 10^{-7}$ |
| Max learning rate | 0.00015 | $3.78 \times 10^{-5}$ |
| Warmup epochs | 14 | $1.30 \times 10^1$ |
| Mode | regression | classification |

**Extended Data Figure 2:**
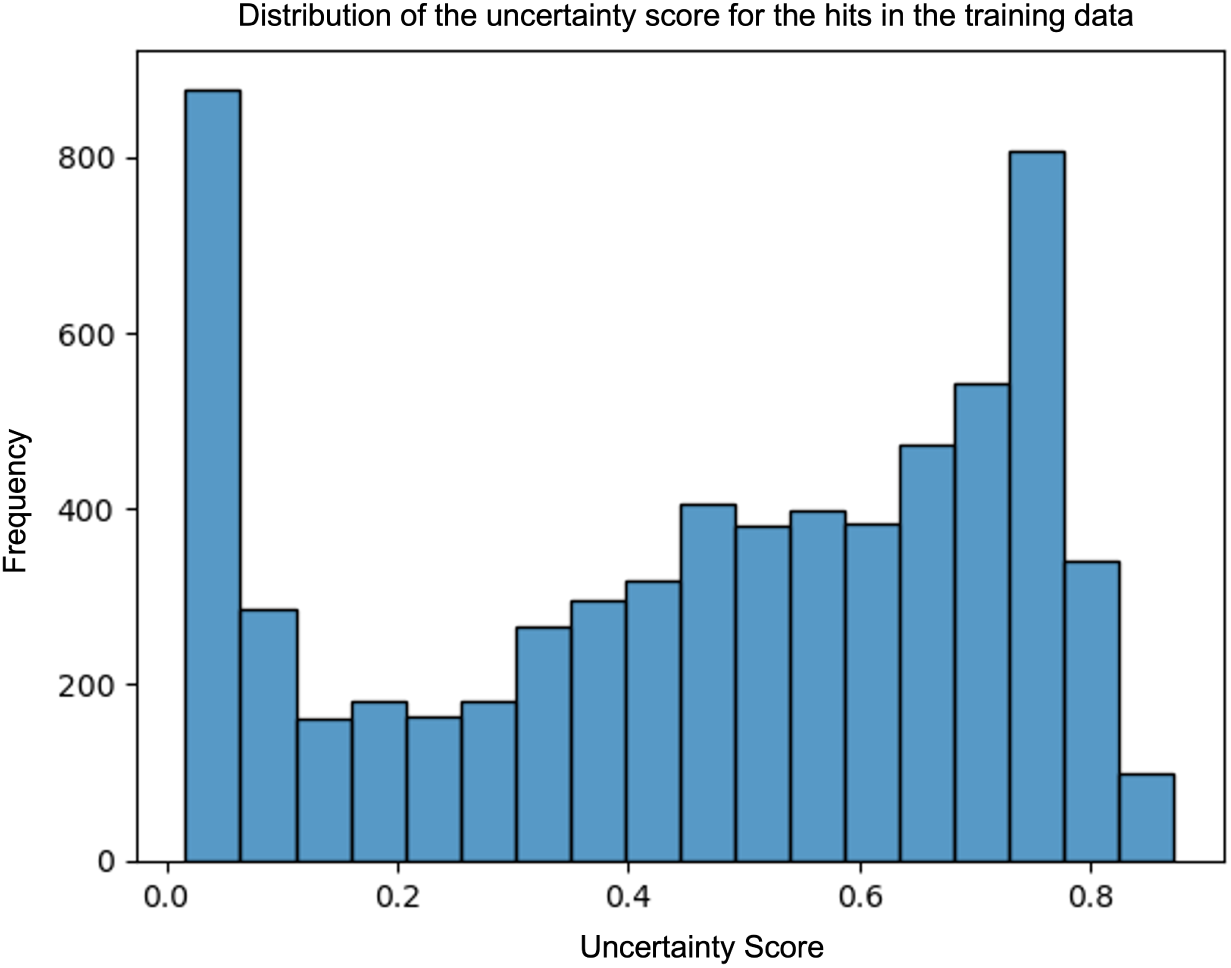
Performance of the XGBoost model on the hits in the training data.

**Extended Data Table 3:** Hyperparameters of the GCNs of the second method (the Chemprop models).

| Hyperparameter | Antibiotic Model | Toxicity Model |
| --- | --- | --- |
| Depth | 2 | 5 |
| Dropout | 0.4 | 0.05 |
| ffn Hidden Size | 1200 | 800 |
| ffn number of Layers | 3 | 3 |
| Hidden Size | 300 | 500 |
| Mode | classification | classification |

**Extended Data Table 4:** Hyperparameters of the XGBoost model used in the first method.

| Hyperparameter | Description | Value |
| --- | --- | --- |
| reg_lambda | L2 Regularization term | 0.1 |
| reg_alpha | L1 Regularization term | 1 |
| learning_rate | Learning rate | 0.1 |
| n_estimators | Number of boosting trees | 800 |
| max_depth | Maximum depth of a tree | 5 |
| gamma | Minimum loss reduction required for splitting | 0 |
| colsample_bytree | Fraction of features randomly sampled for building each tree | 0.3 |

**Extended Data Figure 3:**
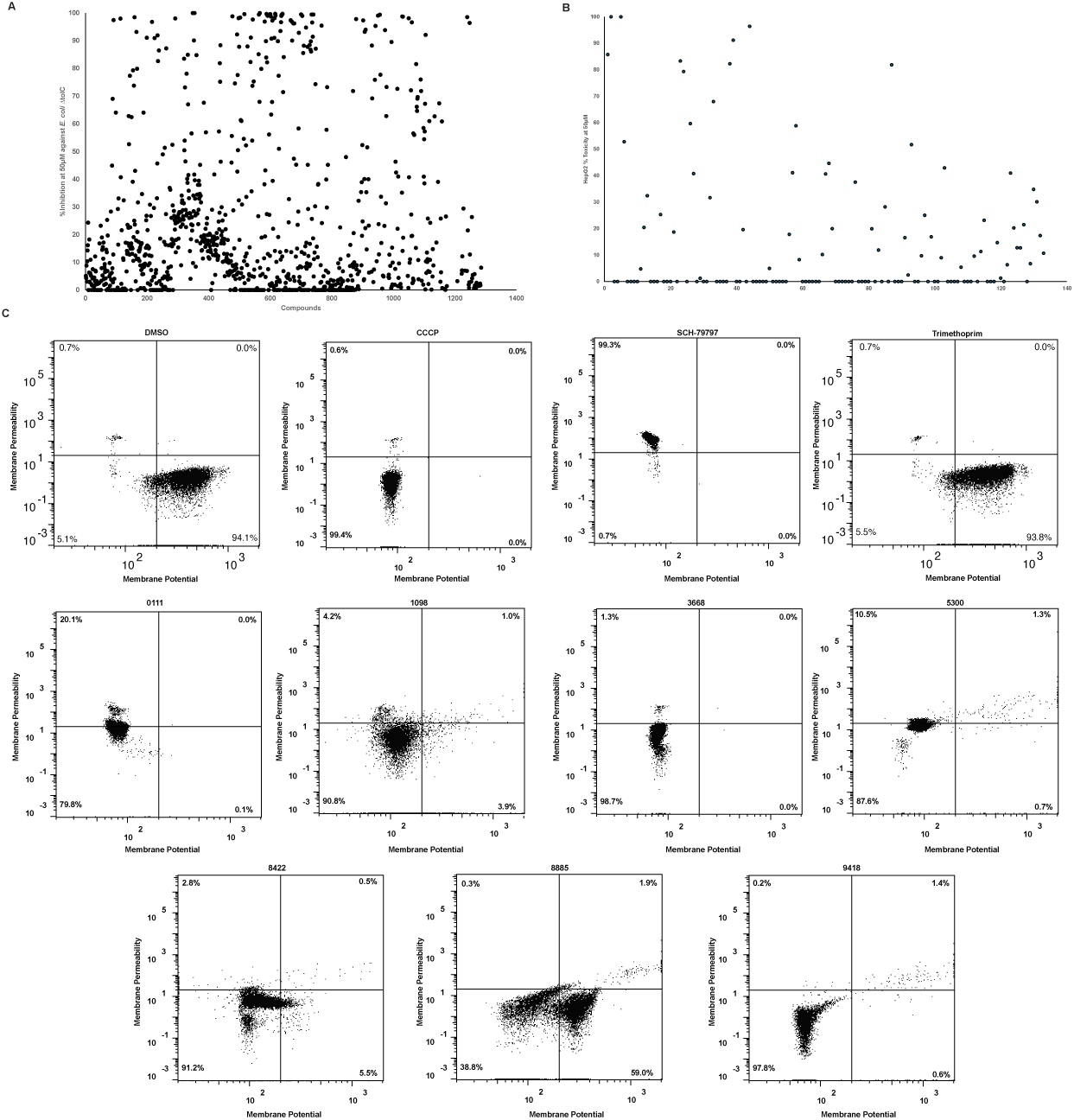
(A) Growth inhibition calculated using optical density at 600nm (OD600)against *E. coli* lptD4213 and *E. coli* ΔtolC, compounds with negative inhibition values are plotted at 0. (B) Percent viability of compounds against HepG2 cells after 24 hours treatment compared to vehicle-only controls. (C) Flow cytometry of active compounds against *E. coli* lptD4213 and stained with TO-PRO-3 and DiOC2(3). Controls for vehicle only (DMSO), CCCP (depolarization), SCH-79797 (permeabilization), and not membrane active (TMP) are also included.

## Notes

### Competing Interest Statement

BEE is on the Scientific Advisory Board for ArrePath, Inc, GSK AI for Cancer, and Freenome; ZG is the Founder of ArrePath, Inc.

## References

1. Murray CJ, Ikuta KS, Sharara F, Swetschinski L, Aguilar GR, Gray A, et al. Global burden of bacterial antimicrobial resistance in 2019: a systematic analysis. The Lancet. 2022;399(10325):629–655.

2. O’Neill J. Antimicrobial resistance: tackling a crisis for the health and wealth of nations. Reviews of Antimicrobial Resistance. 2014;.

3. Göttig S, Gruber TM, Higgins PG, Wachsmuth M, Seifert H, Kempf VA. Detection of pan drug-resistant *Acinetobacter baumannii* in Germany. Journal of Antimicrobial Chemotherapy. 2014;69(9):2578–2579.

4. Nordmann P, Naas T, Poirel L. Global spread of carbapenemase-producing Enterobacteriaceae. Emerging infectious diseases. 2011;17(10):1791.

5. Clardy J, Fischbach MA, Walsh CT. New antibiotics from bacterial natural products. Nature Biotechnology. 2006;24(12):1541–1550.

6. Lewis K. Recover the lost art of drug discovery. Nature. 2012;485(7399):439–440.

7. Ito T, Masubuchi M. Dereplication of microbial extracts and related analytical technologies. The Journal of Antibiotics. 2014;67(5):353–360.

8. Jones S. Permeability rules for antibiotic design. Nature Biotechnology. 2017;35(7):639–639.

9. Chan HCS, Shan H, Dahoun T, Vogel H, Yuan S. Advancing Drug Discovery via Artificial Intelligence. Trends in Pharmacological Sciences. 2019;40:592–604. doi:10.1016/j.tips.2019.06.004.

10. Tommasi R, Brown DG, Walkup GK, Manchester JI, Miller AA. ESKAPEing the labyrinth of antibacterial discovery. Nature reviews Drug discovery. 2015;14(8):529–542.

11. Paul SM, Mytelka DS, Dunwiddie CT, Persinger CC, Munos BH, Lindborg SR, et al. How to improve R&D productivity: the pharmaceutical industry’s grand challenge. Nature reviews Drug discovery. 2010;9(3):203–214.

12. Ramsundar B, Eastman P, Walters P, Pande V. Deep Learning for the Life Sciences: Applying Deep Learning to Genomics, Microscopy, Drug Discovery, and More. O’Reilly; 2019. Available from: https://books.google.com/books?id=tYFKuwEACAAJ.

13. Wu Z, Ramsundar B, Feinberg E, Gomes J, Geniesse C, Pappu AS, et al. MoleculeNet: a benchmark for molecular machine learning. Chemical Science. 2018;9:513–530. doi:10.1039/C7SC02664A.

14. Rahman ASMZ, Liu C, Sturm H, Hogan AM, Davis R, Hu P, et al. A machine learning model trained on a high-throughput antibacterial screen increases the hit rate of drug discovery. PLOS Computational Biology. 2022;18:e1010613. doi:10.1371/journal.pcbi.1010613.

15. Heid E, Greenman KP, Chung Y, Li SC, Graff DE, Vermeire FH, et al. Chemprop: a machine learning package for chemical property prediction. Journal of Chemical Information and Modeling. 2023;64(1):9–17.

16. Yang K, Swanson K, Jin W, Coley C, Eiden P, Gao H, et al. Analyzing Learned Molecular Representations for Property Prediction. Journal of Chemical Information and Modeling. 2019;59:3370–3388. doi:10.1021/acs.jcim.9b00237.

17. Stokes JM, Yang K, Swanson K, Jin W, Cubillos-Ruiz A, Donghia NM, et al. A Deep Learning Approach to Antibiotic Discovery. Cell. 2020;180(4):688–702.e13. 10.1016/j.cell.2020.01.021.

18. Liu G, Catacutan DB, Rathod K, Swanson K, Jin W, Mohammed JC, et al. Deep learning-guided discovery of an antibiotic targeting Acinetobacter baumannii. Nature Chemical Biology. 2023;19(11):1342–1350.

19. Zheng EJ, Valeri JA, Andrews IW, Krishnan A, Bandyopadhyay P, Anahtar MN, et al. Discovery of antibiotics that selectively kill metabolically dormant bacteria. Cell Chemical Biology. 2024;31:712–728.e9. doi:10.1016/j.chembiol.2023.10.026.

20. Wong F, Zheng EJ, Valeri JA, Donghia NM, Anahtar MN, Omori S, et al. Discovery of a structural class of antibiotics with explainable deep learning. Nature. 2023;doi:10.1038/s41586-023-06887-8.

21. Rácz A, Bajusz D, Héberger K. Life beyond the Tanimoto coefficient: similarity measures for interaction fingerprints. Journal of cheminformatics. 2018;10:1–12.

22. Duvenaud DK, Maclaurin D, Iparraguirre J, Bombarell R, Hirzel T, Aspuru-Guzik A, et al. Convolutional networks on graphs for learning molecular fingerprints. Advances in neural information processing systems. 2015;28.

23. Irwin JJ, Tang KG, Young J, Dandarchuluun C, Wong BR, Khurelbaatar M, et al. ZINC20—A Free Ultralarge-Scale Chemical Database for Ligand Discovery. Journal of Chemical Information and Modeling. 2020;60:6065–6073. doi:10.1021/acs.jcim.0c00675.

24. Natividad Ruiz DK, Silhavy TJ. Advances in understanding bacterial outer-membrane biogenesis. Nature Reviews Microbiology. 2006;4:57–66. doi:10.1038/nrmicro1322.

25. Thomas J, Navre M, Rubio A, Coukell A. Shared platform for antibiotic research and knowledge: A collaborative tool to SPARK antibiotic discovery. ACS infectious diseases. 2018;4(11):1536–1539.

26. Nimgaonkar I, Archer NF, Becher I, Shahrad M, LeDesma RA, Mateus A, et al. Isocotoin suppresses hepatitis E virus replication through inhibition of heat shock protein 90. Antiviral Research. 2021;185:104997. doi:10.1016/j.antiviral.2020.104997.

27. Gayvert K, Madhukar N, Elemento O. A Data-Driven Approach to Predicting Successes and Failures of Clinical Trials. Cell Chemical Biology. 2016;23:1294–1301. doi:10.1016/j.chembiol.2016.07.023.

28. Richard AM, Huang R, Waidyanatha S, Shinn P, Collins BJ, Thillainadarajah I, et al. The Tox21 10K Compound Library: Collaborative Chemistry Advancing Toxicology. Chemical Research in Toxicology. 2021;34:189–216. doi:10.1021/acs.chemrestox.0c00264.

29. Kanehisa M, Goto S. KEGG: kyoto encyclopedia of genes and genomes. Nucleic acids research. 2000;28(1):27–30.

30. Lipinski CA. Lead- and drug-like compounds: the rule-of-five revolution. Drug Discovery Today: Technologies. 2004;1:337–341. doi:10.1016/j.ddtec.2004.11.007.

31. Arnott JA, Planey SL. The influence of lipophilicity in drug discovery and design. Expert Opinion Drug Discovery. 2012;10:863–875. doi:10.1517/17460441.2012.714363.

32. Rohde PEB, Selzer P. Fast Calculation of Molecular Polar Surface Area as a Sum of Fragment-Based Contributions and Its Application to the Prediction of Drug Transport Properties. Journal of Medicinal Chemistry. 2000;43:3714–7. doi:10.1021/jm000942e.

33. Travis T Wager PRV Xinjun Hou, Villalobos A. Moving beyond rules: the development of a central nervous system multiparameter optimization (CNS MPO) approach to enable alignment of druglike properties. ACS Chemical Neuroscience. 2010;1:435–49. doi:10.1021/cn100008c.

34. Heald IMSL, Brittelli D. Simple selection criteria for drug-like chemical matter. Journal of Medicinal Chemistry. 2001;44:1841–6. doi:10.1021/jm015507e.

35. Baell JB, Holloway GA. New substructure filters for removal of pan assay interference compounds (PAINS) from screening libraries and for their exclusion in bioassays. Journal of Medicinal Chemistry. 2010;53:2719–40. doi:10.1021/jm901137j.

36. Monica Schenone BKW Vlado Dančík, Clemons PA. Target identification and mechanism of action in chemical biology and drug discovery. Nature Chemical Biology. 2013;9:232–240. 10.1038/nchembio.1199.

37. Steenbergen JN, Alder J, Thorne GM, Tally FP. Daptomycin: a lipopeptide antibiotic for the treatment of serious Gram-positive infections. Journal of antimicrobial Chemotherapy. 2005;55(3):283–288.

38. Velkov T, Roberts KD, Nation RL, Thompson PE, Li J. Pharmacology of polymyxins: New insights into an ‘old ‘class of antibiotics. Future microbiology. 2013;8(6):711–724.

39. Blank J, Deb K. pymoo: Multi-Objective Optimization in Python. IEEE Access. 2020;8:89497–89509.

40. Dice LR. Measures of the amount of ecologic association between species. Ecology. 1945;26(3):297–302.

41. Englert P, Kovács P. Efficient heuristics for maximum common substructure search. Journal of chemical information and modeling. 2015;55(5):941–955.

42. Tversky A. Features of similarity. Psychological review. 1977;84(4):327.

43. Swanson K, Liu G, Catacutan D, Zou J, Stokes J. Generative AI for designing and validating easily synthesizable and structurally novel antibiotics. In: NeurIPS 2023 Generative AI and Biology (GenBio) Workshop; 2023.

44. Zeng X, Xiang H, Yu L, Wang J, Li K, Nussinov R, et al. Accurate prediction of molecular properties and drug targets using a self-supervised image representation learning framework. Nature Machine Intelligence. 2022;4(11):1004–1016.

45. Richter MF, Drown BS, Riley AP, Garcia A, Shirai T, Svec RL, et al. Predictive compound accumulation rules yield a broad-spectrum antibiotic. Nature. 2017;545:299–304. doi:10.1038/nature22308.

46. Geddes EJ, Gugger MK, Garcia A, Chavez MG, Lee MR, Perlmutter SJ, et al. Porin-independent accumulation in Pseudomonas enables antibiotic discovery. Nature. 2023;624:145–153. doi:10.1038/s41586-023-06760-8.

47. Landrum G, Tosco P, Kelley B, Rodriguez R, Cosgrove D, Vianello R, et al.. rdkit/rdkit: 2024_03_1 (Q1 2024) Release; 2024. Available from: 10.5281/zenodo.11102446.

48. Novo D, Perlmutter NG, Hunt RH, Shapiro HM. Accurate flow cytometric membrane potential measurement in bacteria using diethyloxacarbocyanine and a ratiometric technique. Cytometry: The Journal of the International Society for Analytical Cytology. 1999;35(1):55–63.

